# Sc-HistoMorphomics: A General Framework for High-throughput, Multimodal Profiling of Nuclear Morphology in Tissues

**DOI:** 10.64898/2026.11.17.767301

**Authors:** Elaheh Alizadeh, Yue Zhao, Sergii Domanskyi, Dylan Baker, Diane Luo, William F. Flynn, Duygu Ucar, Jeffrey H. Chuang, Elise Courtois, Brian S. White, Nadia Rosenthal, Paul Robson, Ron Korstanje, Sheng Li, J. Matthew Mahoney

## Abstract

Quantitative cellular morphology provides a rich but underutilized window into cell identity and state. We introduce sc-HistoMorphomics, a scalable framework for profiling nuclear morphology in whole-slide H&E images and correlating these measurements with spatial transcriptomics data. Our pipeline combines deep-learning based segmentation with 194 interpretable morphometric features capturing nuclear shape, eosin and hematoxylin texture, and neighbourhood architecture, enabling high-throughput, multimodal characterization of tissue organization. Applied to endometrium, breast cancer tissue, and pancreas, sc-HistoMorphomics revealed conserved high-dimensional feature structure across tissues and reproducible association between morphology and gene expression. Eosin-based texture features were associated with senescence signatures in endometrium, and morphology-based clustering recapitulated broad stromal, lymphocyte, and epithelial tumor cell classes in breast cancer. In pancreas, negative correlations between *CDKN1A* and multiple morphological features were consistent across cell types and donors. Together, these results establish sc-HistoMorphomics as a generalizable and interpretable tool for linking morphology with spatially resolved molecular state across diverse tissues.

## Introduction

Cellular morphology is highly dependent on cell type and function and thus varies substantially within and among biological tissues. Moreover, changes in morphology are often predictive of pathophysiological states, including cancer and aging^1–4^. The morphological state of a cell is a complex function of intrinsic factors, including gene expression, and extrinsic factors, including signaling proteins from neighbouring cells and physical forces acting on the cell boundary. With the advent of spatial omics technologies, it is now possible to molecularly profile cells *in situ* in parallel with classical histology, but spatial omics technologies are low-throughput and expensive, limiting their near-term implementation in clinical applications. Quantitative measures of cell morphology can provide an interpretable phenotypic readout of molecular states identified by spatial transcriptomics. Because H&E images are routinely collected in clinical and research settings, morphology-based features may offer a practical way to relate tissue architecture and cellular phenotype to gene-expression programs measured in more specialized multimodal datasets. Prior studies have shown that image-derived nuclear and histomorphometric features from H&E or related microscopy images can be associated with senescence state, genomic instability, gene-expression programs, prognosis, and clinically used molecular risk scores^2–5,7^. A challenge is that biological processes such as tissue remodeling, cancer progression, and senescence often manifest through subtle, multi-scale alterations in cellular and tissue architecture that are difficult to detect visually and computationally. These changes occur across length scales—from nuclear features to neighbourhood organization—requiring methods that can operate seamlessly from single-cell resolution to entire tissue sections while accounting for cellular heterogeneity.

To address these limitations, we developed sc-HistoMorphomics, a computational framework for quantifying cellular morphology in Hematoxylin and Eosin (H&E)-stained tissue sections at whole-slide scale. Our workflow integrates deep-learning-based segmentation with quantitative morphological features across four distinct categories derived from the nuclear mask, texture-based features of deconvolved eosin and hematoxylin patterns, and neighbourhood features. Each of these captures different and important aspects of cellular and tissue structure. Sc-HistoMorphomics is designed to work at whole-slide scale, using parallel, tile-based processing, to efficiently analyze very large images. This makes it possible to extract high-resolution morphological profiles from hundreds of samples, facilitating integration with additional data modalities.

A key innovation of sc-HistoMorphomics is its ability to integrate morphological features with spatially resolved gene expression data from platforms such as Xenium and Visium, and it can be extended to any dataset that provides spatial coordinates of individual cells (e.g., center of mass in Xenium) or aggregates of cells (e.g., Visium spots). This multimodal framework enables scalable correlation of histological morphology with transcriptomic signatures at scale, supporting the study of complex biological processes and the development of biomarkers from clinically standard histological assessments.

We evaluated sc-HistoMorphomics across diverse tissues and biological states to confirm that it captures meaningful morphological variation at single-cell resolution. When paired with spatial transcriptomics, the framework links histological morphology with gene expression patterns in a scalable and interpretable manner. This positions sc-HistoMorphomics as a generalizable tool for multimodal tissue analysis.

## Results

### A three-component workflow yields interpretable, quantitative morphology features at whole-slide scale

sc-HistoMorphomics was implemented in three main steps (Figure 1). First, we segmented nuclei in images from H&E-stained tissues using the deep learning-based StarDist algorithm^8^. Next, we performed stain separation to isolate the hematoxylin stain, which binds DNA to identify nuclei, from the eosin stain, which binds proteins to identify the cytoplasm and extracellular matrix. Finally, using the segmentation masks and separated stains, we extracted 194 quantitative features representing four major morphometric categories: binary-mask features, eosin-stain features, hematoxylin-stain features, and cell-neighbourhood features (see Methods).

**Figure 1:**
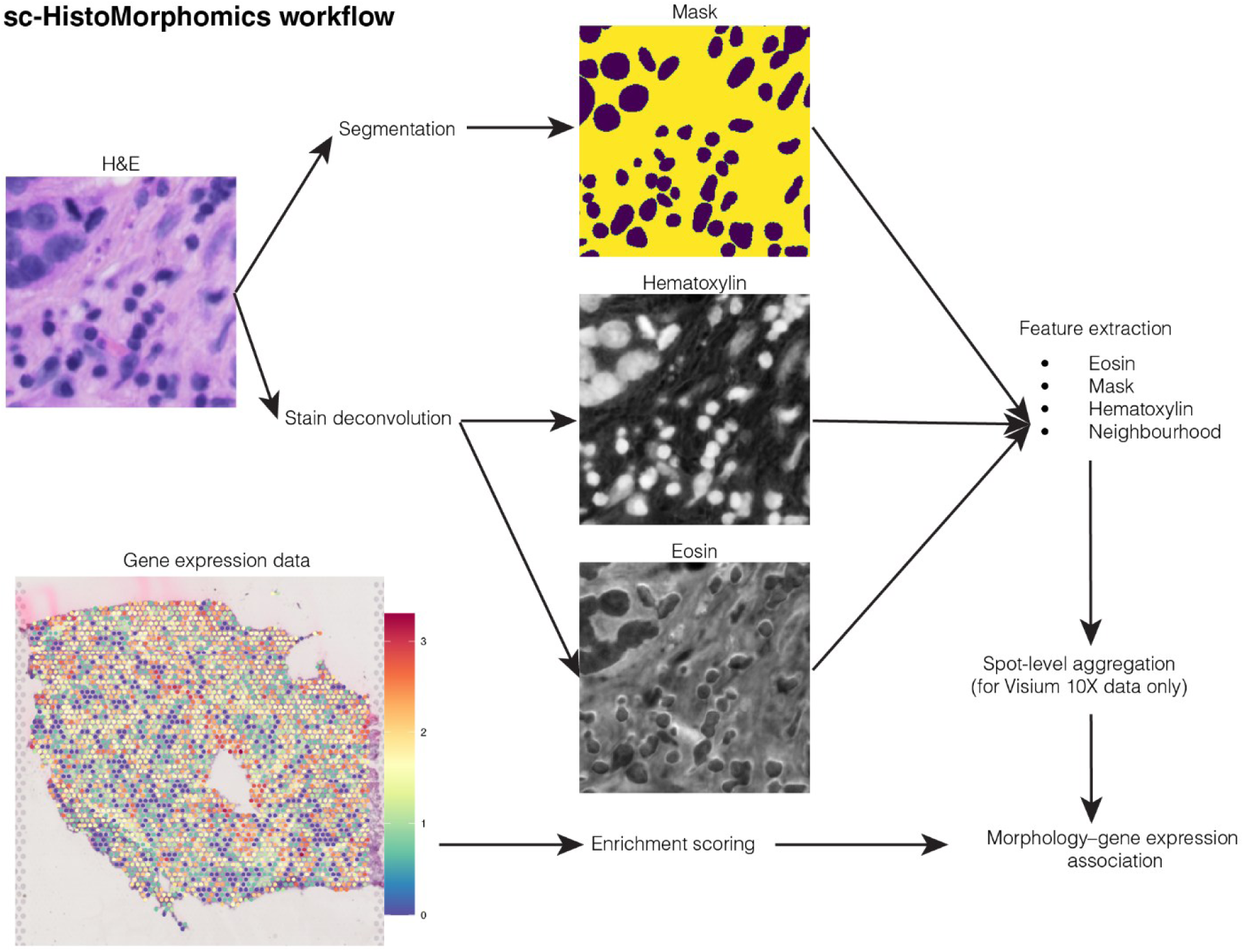
sc-HistoMorphomics framework quantifies 194 nuclear morphological features. Steps for automated morphological quantification of nuclei within H&E images and subsequent correlation with expression-based senescence scores: 1) H&E image is segmented using StarDist to define a nuclei mask. 2) H&E image is split into separate hematoxylin and eosin channels. 3) These two separate images are used to calculate mask, eosin intensity, hematoxylin intensity, and neighbourhood features of each nucleus identified by the mask. 4) Gene set-based senescence scores are calculated for each ST spot (Visium) or cell (Xenium) based on its expression profile. 5) Morphological features are aggregated at the spot level for Visium. 6) Morphological features and senescence scores are correlated at the spot (Visium) or cell (Xenium) level.

Because Visium gene expression is measured at the spot level, we aggregated nuclear morphology features to the same resolution. Specifically, for each spot, we calculated the mean and standard deviation of 194 cell-level morphological features across all nuclei with centroids located within that spot, resulting in 388 spot-level features. These profiles capture both average local morphology and within-spot heterogeneity, enabling direct comparison with spot-level transcriptomic measurements. These features were then correlated with sample-level metadata or with molecular measurements from platforms that pair histological imaging with same-tissue assessment of gene expression profiling, namely Xenium and Visium 10X, to define morphological profiles of cell types and pathophysiologically relevant cell states. We demonstrate this functionality across three tissue contexts: first, morphological clustering in breast cancer tissue and pancreas; next, morphology associated with *CCL5*-expressing cells in breast cancer tissue and *CDKN1A* expression in pancreas; and finally, senescence-associated morphological profiles in endometrium using SenMayo correlations and random forest modeling. The breast cancer and endometrium datasets were used as single-donor proof-of-concept examples, whereas the multi-donor pancreas dataset (14 female donors aged 23–71 and 18 male donors aged 21–54) was used to assess reproducibility across individuals.

### Morphological feature categories are complementary and yield high-dimensional morphological profiles

Given the large number of morphological features, we first reviewed whether they have complementary information and were minimally redundant. To do so, we applied sc-HistoMorphomics to a published H&E image of the endometrium, which was acquired alongside Visium expression data from the same tissue section^9^. Pairwise Pearson correlations among morphological features spanning different major categories—mask, eosin texture, hematoxylin texture, and neighbourhood—were generally low (|r| typically < 0.1), indicating independence of the categories (Figure 2a, Supplementary Figure S1a-b). Within each category, some features showed high correlation (mostly within eosin features with median of 0.25) (Figure 2b). However, overall, correlations remained low within even the same category, demonstrating that this feature set was intrinsically high dimensional, and thus captured a large dynamic range and suggested the capacity to represent diverse, potentially rare morphological states. The variation in pairwise correlations observed even among features within the same broad class is expected because nominally similar descriptors often quantify different spatial aspects of morphology. For example, different texture statistics capture distinct properties of local intensity organization, different moment orders encode different aspects of object geometry, and different radial intensity bins reflect signal distribution at different distances from the nuclear center. As a result, some features within the same class co-vary strongly, whereas others remain only weakly correlated. This behavior is consistent with the intended design of the feature set as a high-content, complementary representation of nuclear morphology rather than a narrowly redundant collection of measurements.

**Figure 2:**
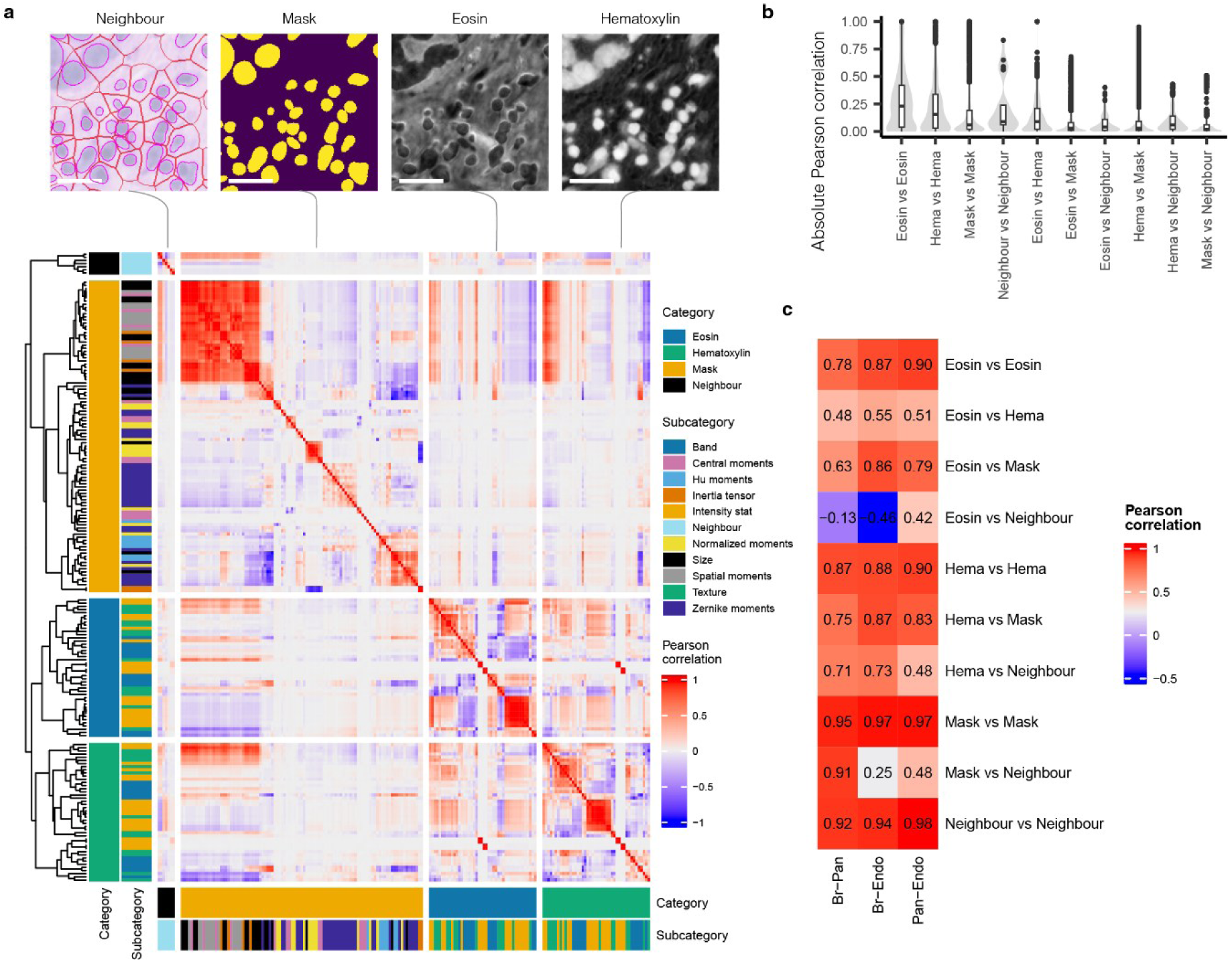
The four major morphological categories are largely independent. **a)** Morphological features are divided into four major categories that are calculated based on a binary mask of nuclei, intensity of the eosin stain within nuclei, intensity of the hematoxylin stain within nuclei, and the neighbouring density of each nucleus. Scale bar, 20 µm. The low correlation coefficient between features shows that these major categories are largely independent and capture different aspects of nuclei morphology. b) Correlation of features within each category is small. c) Correlation of feature correlation across three tissues shows that the relation between features is conserved across the tissues.

To quantify redundancy more directly, we calculated the fraction of feature pairs exceeding high absolute Pearson correlation thresholds. Across all feature pairs, only 2.87% exceeded ∣r∣>0.8, and only 1.44% exceeded ∣r∣>0.9, indicating that most features were not highly redundant. We also summarized absolute pairwise correlations by feature-category pair (Supplementary Table 1). Within-category feature pairs showed higher correlations than between-category pairs, as expected, but median absolute correlations remained modest for many category pairs, supporting the interpretation that the feature set captures complementary aspects of morphology.

To further evaluate dimensionality, we performed PCA on the z-scored morphology feature matrix. The variance explained by individual principal components decreased gradually rather than being concentrated in only a few components. Forty-three principal components were required to explain 90% of the variance, and 60 principal components were required to explain 95% of the variance. (Supplementary Figure S2)

Low pairwise correlation among many features does not imply that the feature space lacks informative structure; rather, it indicates that the measured descriptors capture complementary aspects of morphology. Together, these results indicate that the 194-feature representation is high-dimensional and only partially redundant, capturing multiple modes of variation across nuclear shape, eosin texture, hematoxylin texture, and neighbourhood organization.

We also quantified effective dimensionality using the spectral-entropy “effective rank”^10^ of the singular value spectrum after z-scoring features. The effective rank was 95 for endometrium and 93 for both breast and pancreas, indicating that the variance is spread across ∼93–95 independent patterns, demonstrating relatively low feature redundancy and confirming that the datasets preserve substantial high-dimensional structure.

We next asked whether the relationships among morphology features were preserved across tissues by comparing, for each feature-category pair, the correlation structure observed in breast, endometrium, and pancreas (Figure 2c). Overall, within-category correlation patterns were highly conserved: pairs such as mask–mask, neighbour–neighbour, hematoxylin–hematoxylin, and eosin–eosin showed strong cross-tissue similarity (r ≈ 0.85–0.98). Cross-category interactions involving mask or hematoxylin were also broadly consistent across tissues (e.g., hematoxylin–mask: 0.75–0.87; eosin–mask: 0.63–0.86). In contrast, category pairs involving neighbour exhibited noticeably greater variability. For example, mask–neighbour correlations ranged widely across tissues (0.25–0.91), and eosin–neighbour even changed sign between tissue pairs. This increased variability was expected because the neighbour category contains far fewer features (n = 7) than the other categories, resulting in substantially fewer feature-pair correlations contributing to these comparisons and therefore noisier estimates. Taken together, these results indicated that most aspects of the morphological correlation structure are preserved across tissues, whereas relationships involving neighbour features appeared more tissue-dependent and less stable, reflecting both biological differences in tissue architecture and the smaller feature set contributing to those comparisons.

### Morphological features are robust to segmentation algorithm choice

Because all downstream morphological measurements depend on upstream segmentation, we next assessed the sensitivity of sc-HistoMorphomics to segmentation algorithm choice. We compared nuclei segmented from the same breast cancer H&E image using StarDist and Cellpose, matched nuclei across methods by centroid proximity, and then compared corresponding morphological features using both Pearson and Spearman correlation. StarDist processed the image substantially faster than Cellpose (∼10 min versus ∼7.5 h, respectively, using the same computational resources). Across feature categories, hematoxylin and eosin texture features showed the strongest reproducibility, with most correlations concentrated at high values and many Spearman correlations exceeding 0.7. By contrast, mask-derived shape features showed broader agreement distributions, with the weakest concordance observed primarily for Zernike moment features, which are expected to be more sensitive to subtle differences in nuclear boundary definition; neighbourhood features were the most variable overall. These results indicate that the most biologically informative texture-based features are robust to the choice of segmentation method, whereas boundary-sensitive mask and neighbourhood measurements are more sensitive to local segmentation differences. Importantly, sc-HistoMorphomics is modular with respect to segmentation: users can provide nuclear masks generated by StarDist, Cellpose, or other segmentation algorithms as input to the downstream feature-extraction pipeline. Representative segmentation overlays, feature correlations, and full feature-wise statistics are shown in Supplementary Figure S3 and Supplementary Table S2.

### Morphology-based clustering recovers broad cell classes in breast cancer tissue

To evaluate sc-HistoMorphomics at true single-cell resolution, we applied the framework to a published Xenium single-cell spatial transcriptomics dataset of a breast cancer tissue section^11^. Xenium provides subcellular-resolution transcript localization and cell-level molecular annotations, enabling direct comparison between single-cell morphology and gene-expression-derived cell identities. We therefore quantified morphological features of individual nuclei in the paired H&E image without subsequent aggregation. Unsupervised clustering of these features revealed three dominant morphological classes (Figure 3a) corresponding to the three published expression-based clusters of stromal, lymphocyte, and epithelial tumor cells (Figure 3b). To quantitatively assess this correspondence, we compared morphology-based clusters with independently defined gene-expression-derived cell type annotations using a cluster–cell type concordance heatmap, where each entry represents the fractional overlap of an expression-defined cell type within a morphology-derived cluster. The morphology-based clusters showed clear enrichment for broad expression-defined cell classes (Figure 3c-d). This highlighted the cell type-specificity of morphological profiles, including stromal cells, which, as in the endometrium, have been associated with senescence in breast cancer^11–13^. We also evaluated whether morphology-based clustering resolved finer transcriptionally defined subtypes within major cell classes. These more closely related stromal and lymphoid subpopulations showed limited separability using H&E-derived morphology alone, indicating that the strongest discriminative signal occurs at the level of broad cell classes and morphology-associated states rather than all molecular subtypes. This analysis showed that morphology-based clusters were enriched for broad cell classes, while not implying one-to-one classification accuracy for all transcriptionally defined subtypes.

**Figure 3:**
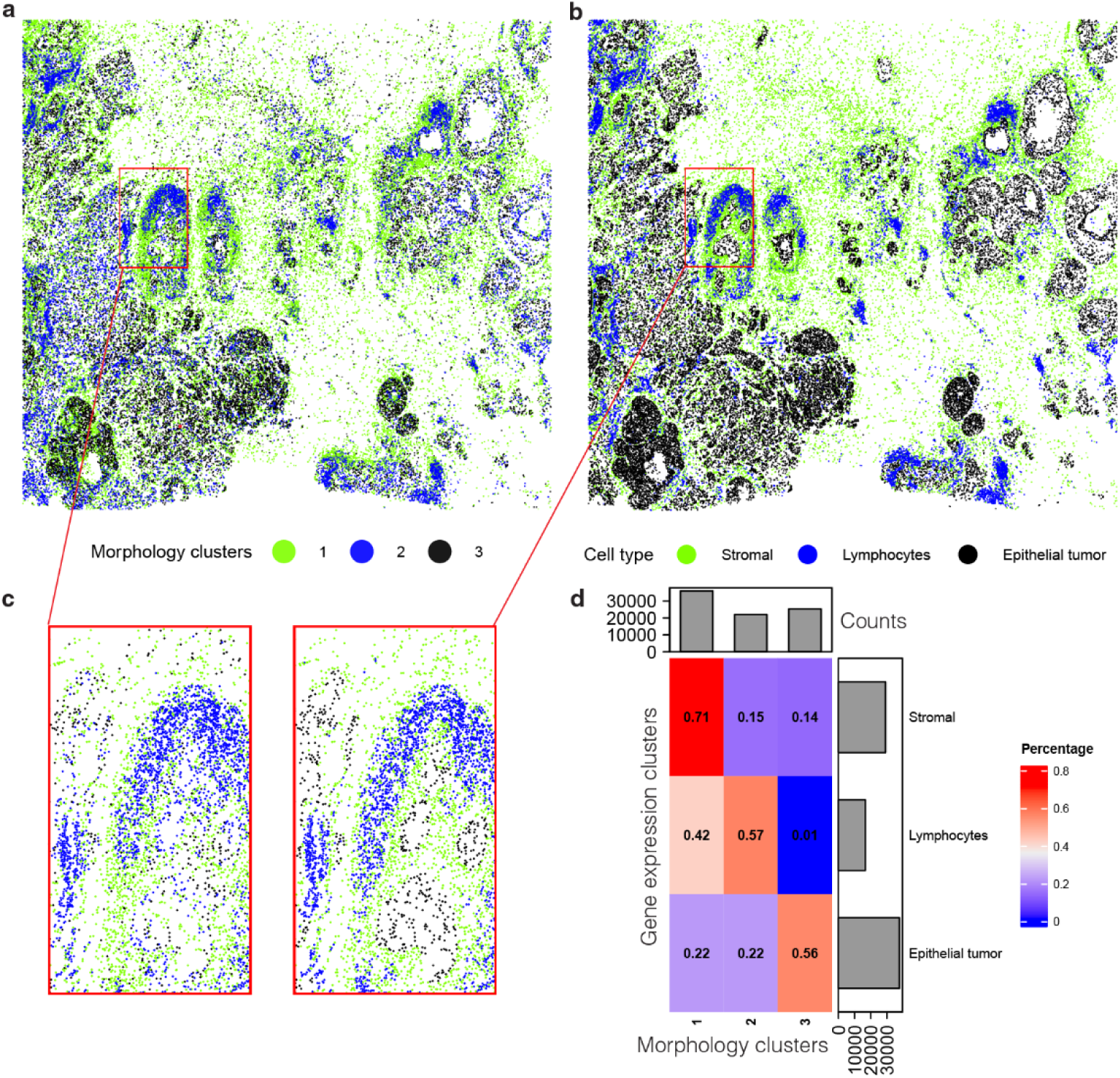
Morphological features of nuclei capture cell type heterogeneity in breast cancer tissue. **a)** Morphological feature clusters derived from Xenium-paired H&E image. b) Published cell types derived from Xenium gene expression data. c) Inset images illustrate examples of morphology-based clusters and corresponding expression-derived cell-type annotations between the three morphological feature-based clusters and gene expression-based cell types. d) Heatmap shows fractional overlap between morphology-based clusters and gene-expression-derived cell type annotations. Values indicate enrichment of each expression-defined cell type within each morphology-derived cluster and should be interpreted as concordance rather than supervised classification accuracy.

### Morphology-based clustering shows partial concordance with pancreatic cell annotations

Having established that sc-HistoMorphomics can recover broad cell classes and within-cell-type molecular heterogeneity in a single-cell Xenium breast cancer dataset, we next asked whether morphology–gene expression relationships were reproducible across individuals. To do so, we analyzed a multi-donor human pancreas Xenium dataset comprising 32 tissue samples from 14 female donors aged 23–71 years and 18 male donors aged 21–54 years^14^. The Xenium panel included 300 genes, including *CDKN1A* and 37 additional senescence-related genes, enabling assessment of morphology-associated transcriptional variation across donors and pancreatic cell types.

Morphology-based clusters showed partial concordance with gene-expression-derived cell type annotations, with strongest enrichment observed for broader cell classes(Figure 4; see Methods). This correspondence was weaker than in the breast cancer dataset, consistent with the greater cellular diversity of pancreas tissue, the subtler morphological differences among some pancreatic cell populations, and increased regional heterogeneity relative to the breast cancer example.

**Figure 4:**
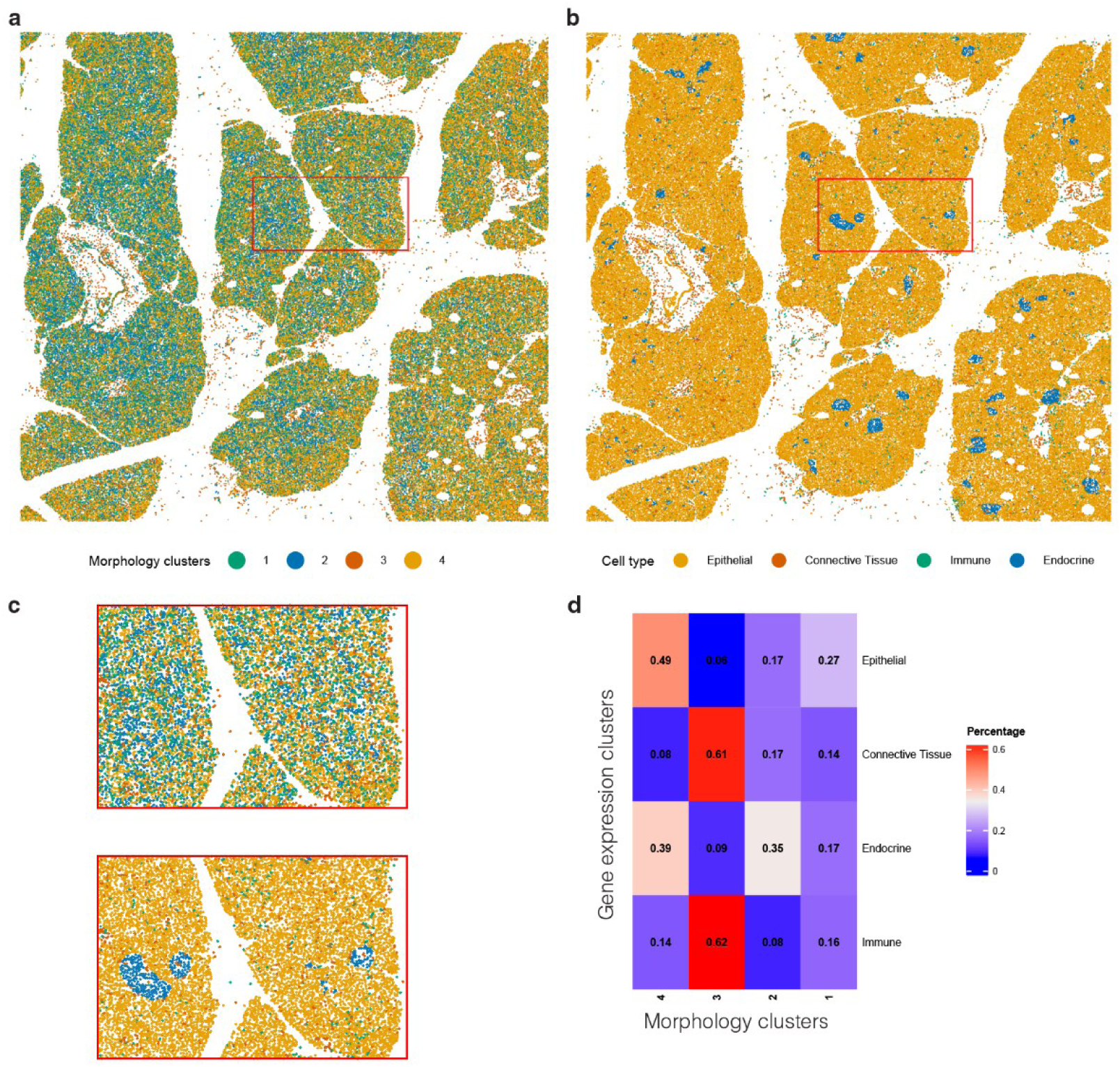
Morphology-based clustering in pancreas tissue shows partial concordance with expression-derived cell-type annotations. **a)** Morphological feature clusters derived from Xenium-paired H&E image. b) Published cell types derived from gene expression of Xenium data. c) Inset images emphasize the high consistency between the four morphological feature-based clusters and gene expression-based cell types. d) Heatmap shows fractional overlap between morphology-based clusters and gene-expression-derived pancreatic cell type annotations. Concordance is strongest for broad cell classes.

### Morphological features capture *CCL5*-associated heterogeneity within stromal cells

We next investigated whether morphological features were associated with expression of cancer-relevant genes. We focused on *CCL5*, a chemokine included in the Xenium probe set that is implicated in immune-cell recruitment, tumor–immune interactions, *SASP*-related inflammatory signaling, and breast cancer progression. For these reasons, we included *CCL5* as a senescence- and inflammation-associated marker in this dataset^15,16^. Within the stroma cells, the highest correlated features with *CCL5* expression were the percentage of cells touching (neighbourhood feature), major axis length (mask feature), and hematoxylin distribution in the boundary of nuclei and the minimum intensity of eosin (intensity-based features; Figure 5a). These representative features make it clear that the *CCL5*-associated signal is distributed across multiple interpretable feature classes, spanning cellular geometry, stain texture, and local tissue context.

**Figure 5:**
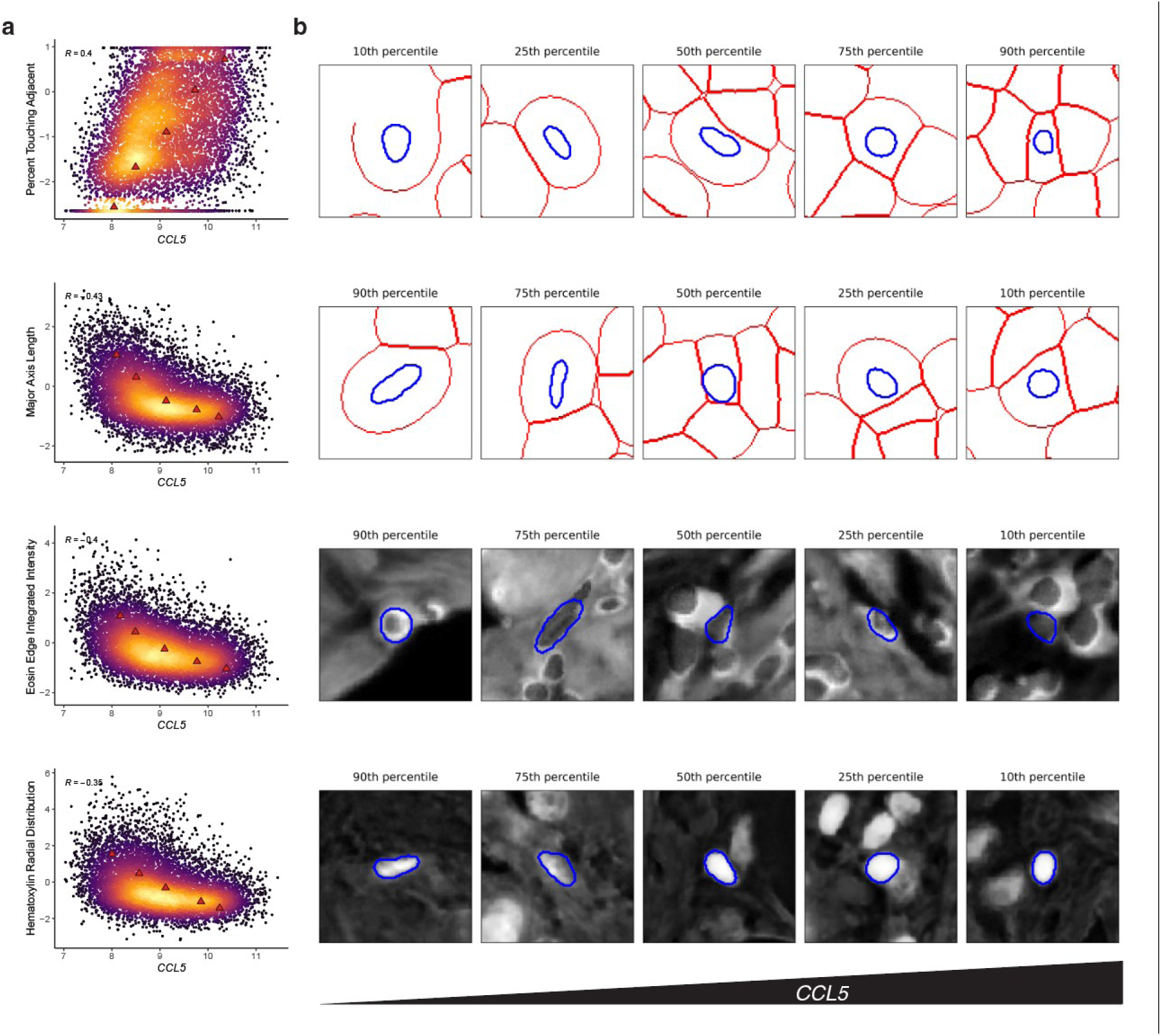
*CCL5* expression correlated with morphological features in breast cancer tissue in stromal cells. a) *CCL5* expression correlation against representative features having the strongest association within each major category. All correlations were significant (P < 2.2 × 10⁻¹⁶). b) Representative images of nuclei at increasing levels of *CCL5* expression and associated quantiles of morphological features from a).

To determine whether these associations reflected molecular variation within a shared cell type, rather than broad stromal versus non-stromal differences, we restricted the analysis to stromal cells and compared *CCL5*-expressing and non-expressing stromal cells for representative morphology features. Several features differed significantly between the two groups, including the neighbourhood feature percent touching adjacent, major axis length, eosin edge integrated intensity, and hematoxylin radial distribution (Supplementary Figure 4). For example, major axis length was lower in *CCL5*-expressing stromal cells than in non-expressing stromal cells (Wilcoxon rank-sum test, p < 2.2 × 10⁻¹⁶). Continuous correlations with *CCL5* expression within stromal cells further supported these within-cell-type relationships. Because binary expressor/non-expressor comparisons and continuous expression correlations address related but distinct questions, they are not necessarily expected to show identical directionality for every feature, particularly for neighbourhood descriptors. Together, these results indicate that sc-HistoMorphomics captures transcriptionally defined heterogeneity within stromal cells, rather than merely broad cell-type identity. Cells with higher *CCL5* expression were densely packed with rounded nuclei (first two rows; Figure 5a-b). In addition, the outermost radial nuclear intensity decreased with increasing *CCL5* expression (third row; Figure 5a-b). Similarly, the integrated eosin edge intensity of the nuclei decreased with increasing *CCL5* expression (fourth row; Figure 5a-b). Together, these features showed that *CCL5* expression, a *SASP* gene that is known to drive cancer progression^17,18^, was strongly correlated to both intracellular and neighbourhood-level morphological traits, demonstrating that sc-HistoMorphomics identified pathophysiologically relevant morphological variation within this breast cancer tissue image.

### Morphological features of single nuclei consistently correlate with *CDKN1A* expression across pancreas donors

Cellular senescence is a multifaceted biological process characterized by stable cell-cycle arrest, altered tissue function, and activation of stress-response and inflammatory programs. Several RNA and protein biomarkers, including *CDKN1A* encoding *p21*, *CDKN2A*-related markers, and components of the senescence-associated secretory phenotype (*SASP*), are well-established and widely used indicators of senescence, although none is fully specific or exhaustive on its own^19^. Although senescence in vitro has been associated with morphological changes, the extent to which senescence-associated transcriptional states are reflected in tissue morphology in vivo remains incompletely understood. We therefore used sc-HistoMorphomics to test whether H&E-derived morphology is associated with transcriptomically defined senescence-enriched states across pancreas and endometrium datasets^20^

Given the theorized importance of cellular senescence in aging pancreas^21^, we correlated morphological features with *CDKN1A* expression (library-size–normalized). We observed high negative correlations between *CDKN1A* and cell area, aspect ratio, and entropy of hematoxylin stain (a texture feature; Figure 6a). These recurrent features were the most reproducible morphology correlates of *CDKN1A* across donors and cell types, indicating that both nuclear geometry and chromatin-associated texture contribute to the senescence-associated morphology signal in pancreas, indicating that the morphological features reflected shared, cross cell type characteristics of senescence (Figure 6B). Figure 6b presents the similar relationship between *CDKN1A* and cell area across cell types as an example. We validated these negative correlations in a separate “head” section of the pancreas from the same patient, as well as within head and tail sections from a second patient (Supplementary Figure 5). Together, these findings demonstrate that the relationship between senescence-associated gene expression and morphological features is conserved across different cell types and patient samples. This suggests that morphology captures transcriptomically defined *CDKN1A*-associated states that are consistent with senescence-associated biology. We also evaluated alternative normalization strategies (Supplementary Figure 6) and observed that correlations with *CDKN1A* remained consistent across methods.

**Figure 6:**
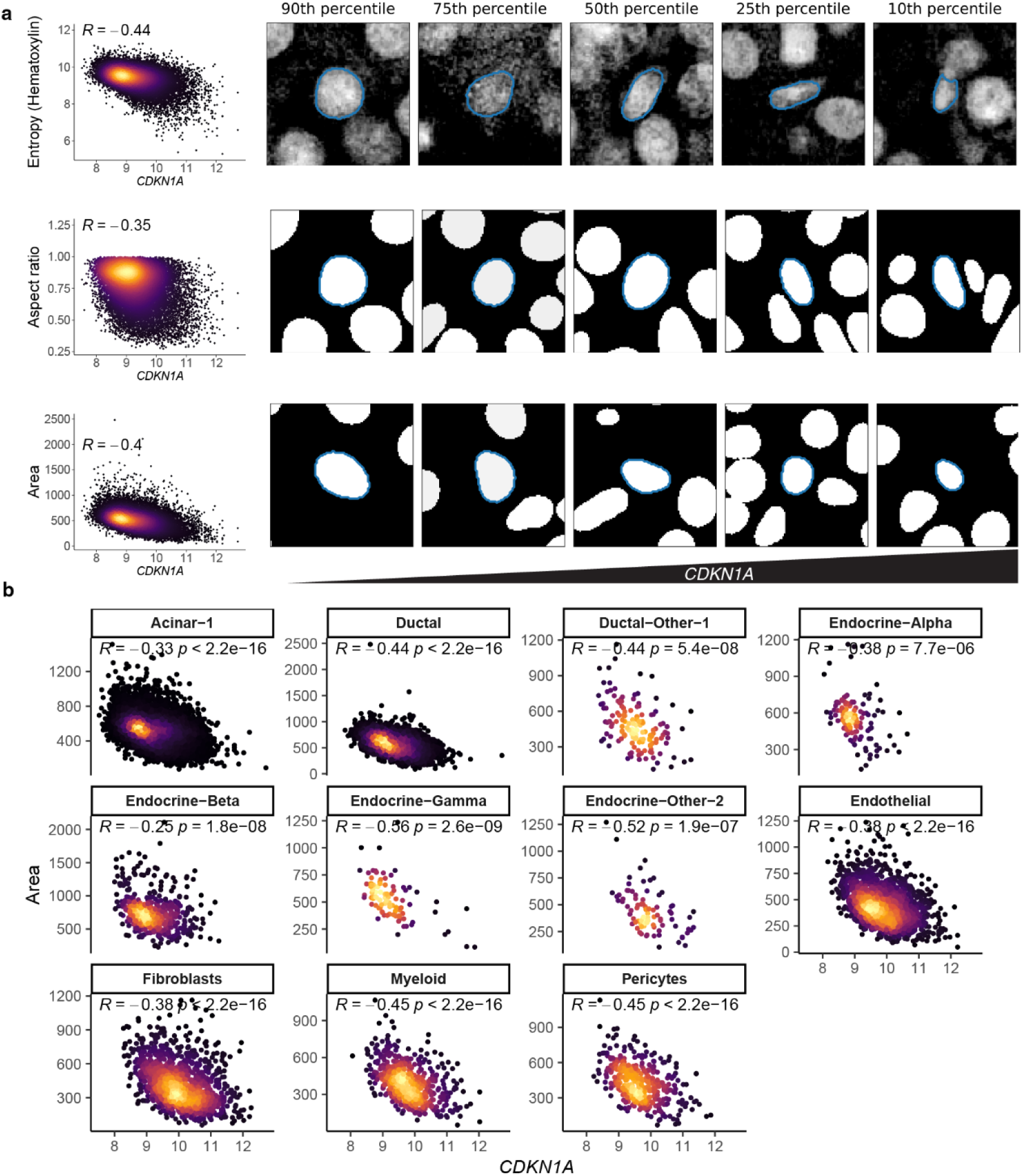
Nuclei shape and texture features correlate with senescence-associated *CDKN1A* in pancreas. **a)** Correlations between *CDKN1A* expression (encoding senescence marker *p21*) and morphological features having largest magnitude correlations: cell area, aspect ratio, and hematoxylin stain entropy. Representative images of nuclei at levels of morphological feature and *CDKN1A* depicted on the left. b) Cell type-specific correlations between *CDKN1A* and area. All gene-expression values reflect library-size–normalized counts.

To determine whether morphology associations extended beyond CDKN1A, we examined morphology correlations across the full 300-gene Xenium panel, including canonical senescence-associated genes and genes related to DNA-damage response and inflammatory signaling. Several senescence-associated genes, including CDKN2A, CDKN1C, TIMP1, and TRIAP1, showed moderate morphology associations (maximum |ρ| ≈ 0.3-0.6). Across these genes, the strongest positive correlations were consistently observed for texture-related features, particularly Angular Second Moment measurements, whereas the strongest negative correlations occurred with cell size and geometry features such as maximum radius, Feret diameter, and inertia tensor eigenvalues. Similar morphology associations were observed across many non-senescence genes in the panel, suggesting that senescence-associated genes participate in a broader morphology program rather than defining a unique morphology signature.

### Eosin texture features identify senescence-associated morphology in endometrium

#### Endometrial tissue architecture provides a context for senescence-associated morphology

We next analyzed Visium spatial transcriptomics of the endometrium alongside an H&E image acquired from the same tissue section^9^. The endometrium, which lines the uterus, is a tissue composed of epithelial glands surrounded by stroma. The stroma is particularly relevant because it is enriched for senescent cells, as demonstrated by the abundant presence and spatial distribution of *p16*-positive cells in the endometrial stromal compartment during the implantation window^22^. This structural organization - gland surrounded by senescence enriched stromal tissue - provides a clear morphological context for evaluating senescence associated morphological features and a useful morphological context for evaluating senescence-associated features.

#### SenMayo scores identify stromal-enriched senescence-associated regions

To identify senescence-enriched spots for subsequent morphological characterization, we calculated expression-based senescence scores for each spot using the SenMayo gene sets. The SenMayo gene set comprises 125 senescence-associated genes organized into different functional clusters: metalloproteases, cytokine/chemokine, growth factors, intercellular signal molecule, protease Inhibitors, and transmembrane signal receptor. We also computed spot-level enrichments of endometrium-relevant cell types using curated gene sets (see Methods). Cell type enrichments were significantly correlated with well-established marker genes, including *EPCAM* (epithelial), *VIM* (stromal) (Supplementary Figure 7). Spatial profiling studies of endometrium show that senescent cells are located within the stroma^22^. As anticipated, spots with a high SenMayo score were enriched for stromal cells and depleted for epithelial cells in the endometrium (Supplementary Figure 8). Notably, the spots with high senescent-associated expression were dispersed throughout the tissue (Supplementary Figure 9). The enrichments related to protease inhibitor, metalloprotease, and intracellular signal had the most heterogeneous distribution across the tissue (Supplementary Figure 9). These three scores were also correlated with each other (Supplementary Figure 10). As each subcategory represents a distinct, non-overlapping set of genes, their correlation suggests they capture related but biologically distinct senescence-associated processes.

#### Eosin texture features show the strongest association with SenMayo

We next generated morphological features for each nucleus, calculated spot-level morphological features by computing mean and standard deviation across nuclei in each spot (388 features total), and correlated spot-level morphological features with spot-level senescence scores (Figure 7a, see Methods). Across the four major morphometric categories, eosin-derived texture features showed the strongest correlation with senescence-related gene expression. When examining absolute Pearson correlations between spot-level mean features and the SenMayo score, eosin features exhibited both the highest median effect size (median |r| = 0.238) and the broadest interquartile range (IQR = 0.201), with individual features reaching correlations as high as |r| = 0.364. Hematoxylin texture features showed more moderate associations (median |r| = 0.0667, IQR = 0.144, max |r| = 0.290), while mask-based nuclear shape measures and neighbourhood features displayed weaker but consistent relationships (mask: median |r| = 0.0525; neighbourhood: median |r| = 0.0618). These patterns highlighted that subtle eosin texture variations, which capture cytoplasmic and extracellular matrix organization, encode substantially more senescence-related information than nuclear shape alone. Collectively, these results underscored the value of texture-based features—particularly those derived from eosin staining—as sensitive morphological markers of senescence-associated tissue alterations (Figure 7b). Among all individual morphology features, the strongest association with SenMayo was the mean texture correlation of the eosin stain, making it the most informative single feature in the endometrium analysis (Figure 7). This feature is calculated based on the gray-level co-occurrence matrix (see Methods) and measures the linear relationship between neighbouring pixel intensities in an image. A high texture correlation (near 1) reflects a smooth, uniform texture with strongly predictable intensity patterns, whereas a low correlation (near 0) indicates a heterogeneous or noisy texture with rapid, less predictable intensity changes. We observed that cells with uniform texture have low SenMayo score and cells with random eosin texture have higher SenMayo score.

**Figure 7:**
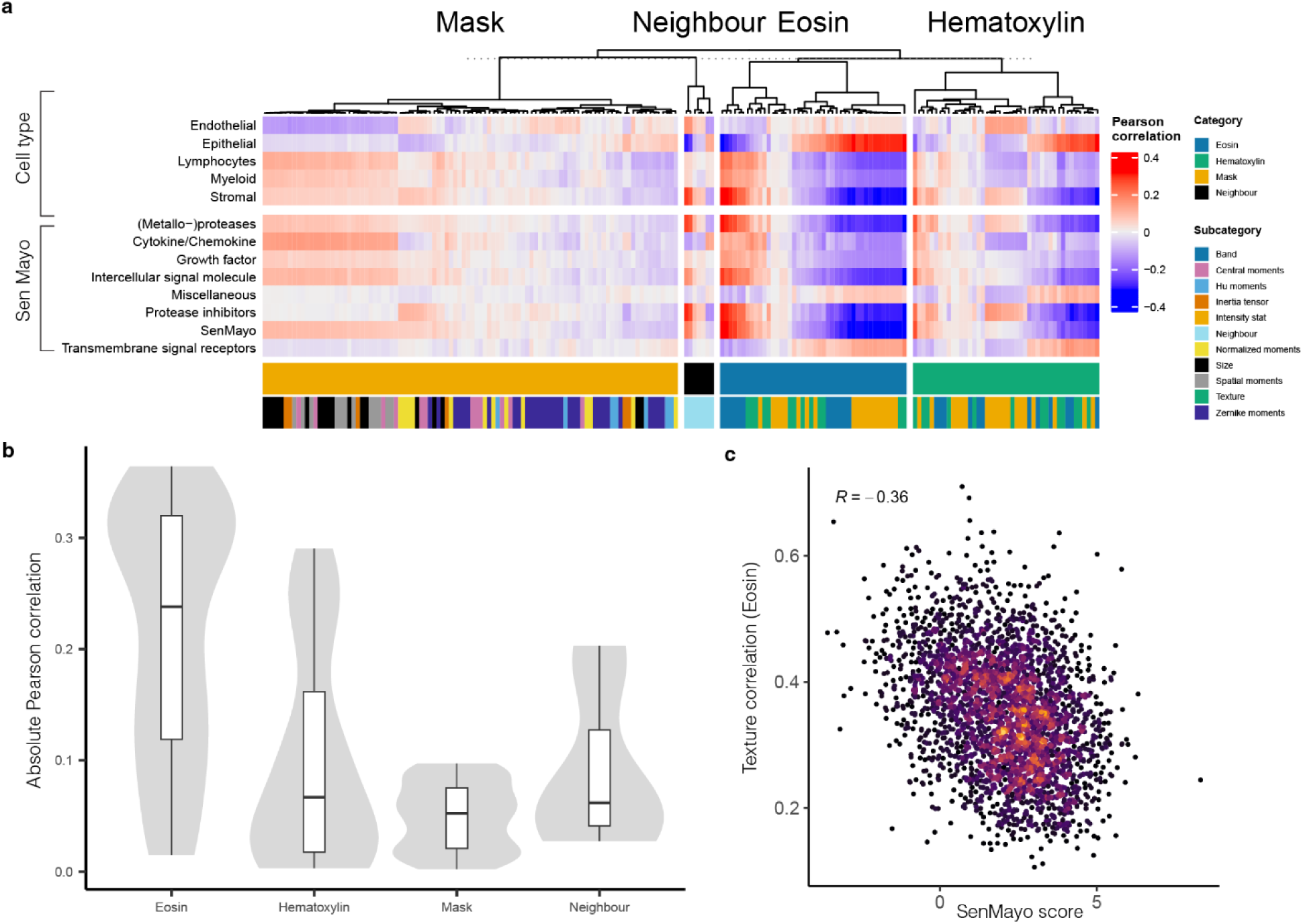
Average eosin features within each spot are morphological markers of senescence in the endometrium. **a)** Pearson correlation between average morphological features within a spot and either spot-level SenMayo scores or cell type enrichments. b) Features related to average values of eosin features per spot show largest associations with SenMayo score. c) Correlation of the SenMayo score with the top morphological feature, eosin texture correlation.

Because Visium measurements are made at the spot level, and SenMayo-enriched spots in the endometrium were also enriched for stromal cells, we next asked whether the top morphology–SenMayo association was explained solely by the cellular composition of each spot. We therefore performed multivariate linear modeling that adjusted for stromal and epithelial composition and found that the top eosin texture feature (mean texture correlation) remained significantly associated with SenMayo (β = −1.63, p = 1.21 × 10⁻¹⁰). Moreover, adding this feature significantly improved model fit relative to a composition-only model (nested model comparison, F = 41.74, p = 1.21 × 10⁻¹⁰). These results indicate that the eosin texture signal is not explained solely by stromal versus epithelial composition, but captures additional senescence-associated transcriptional variation at the spot level.

To assess whether local morphological variability within each spot also relates to senescence, we examined the standard deviation of each feature across each spot. This provides a complementary measure of intra-spot heterogeneity, capturing whether senescent regions differ not only in their average texture or intensity but also in how variable these properties are among adjacent cells. The standard deviations showed a similar pattern to the means: texture features derived from the eosin stain remained among the most strongly correlated with the SenMayo score (Supplementary Figure 11).These results demonstrated that putative senescent spots were morphologically distinct from surrounding tissue based on subtle texture-based features that quantify how smoothly or irregularly eosin stains vary across small spatial neighbourhoods—both in terms of average texture and the heterogeneity of those textures within each spot.

#### Multivariate morphology profiles improve prediction of SenMayo scores

Given that spatial correlations between the SenMayo score and several morphological features remained significant after correction for multiple testing, we next asked whether an optimized combination of them would more strongly associate with senescence than any individual feature alone. While the univariate analyses highlight informative associations (Figure 7), they inherently consider each feature in isolation and therefore cannot capture nonlinear relationships, interactions between features, or composite texture patterns that may collectively reflect senescence-associated features. To address this, we trained a random forest model to predict the SenMayo score as a function of the full 388-feature morphological profile. Our out of the bag performance achieved Pearson correlation performance of *R*=0.48 between predicted and observed SenMayo scores (Figure 8a, b). This performance was stable under 5-fold cross-validation, where each fold produced similar correlation values. This performance was higher than the correlation of any individual feature with SenMayo score (0.37, Figure 7c), indicating that the model leverages information across features to improve prediction accuracy. The feature importance rankings for the model showed that the mean eosin-intensity-based features had the highest influence on overall model predictions (Figure 8c). The most important individual feature was the mean texture correlation of the eosin stain, consistent with the above univariate results in which it achieved the highest magnitude correlation with the SenMayo score, indicating that eosin-derived texture is a recurrent driver of the senescence-associated morphology signal (Figure 7c).

**Figure 8:**
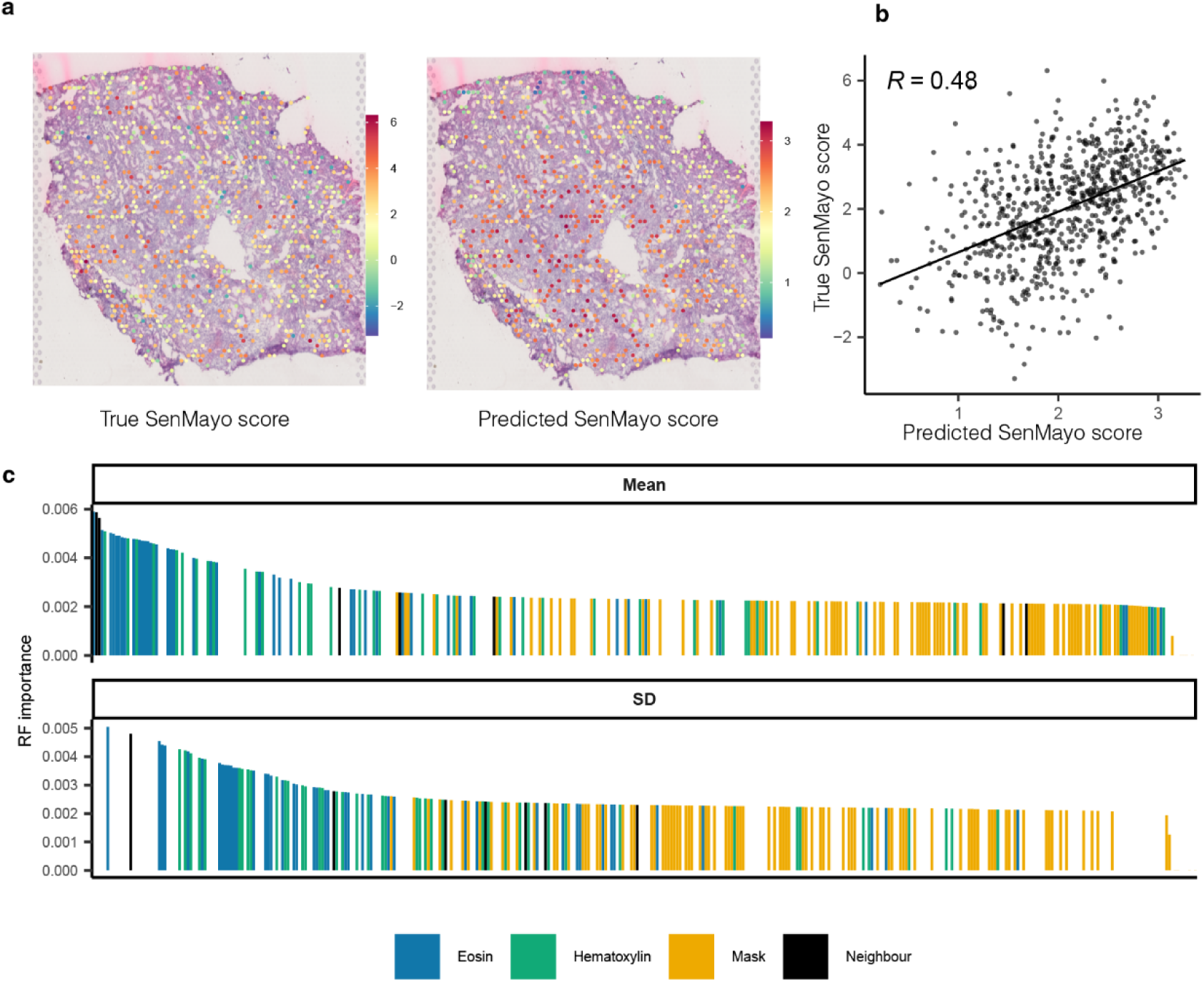
Morphological features predict senescence score in the endometrium. **a)** True (left) and random forest prediction (right) of SenMayo score. b) Correlation of true (y axis) and predicted (x axis) SenMayo scores. Random forest predictions evaluated on a subset of held-out spots within the tissue disjoint from those spots used for training. c) Variable importance of features within trained random forest model, highlighted by morphological category, and separated according to the mean (top) or standard deviation (SD; bottom) of nuclei features within a spot. Mean eosin features are those most important to the model.

## Discussion

Our pipeline enables *in situ* exploration of relationships *between* histological morphology and transcriptomic signatures at scale, providing a framework to study complex cellular phenotypes in tissues. This multimodal capability of the pipeline is particularly valuable for complex biological processes such as cellular senescence, where rare cell populations and heterogeneous phenotypes require both high-resolution imaging and transcriptomic data. The scalability of our approach enables tissue-wide surveys capable of capturing rare cell types; notably, processing ∼90,000 cells required 15 CPU cores, ∼64 GB of memory, and a 4 wall-clock time. Together, our results establish sc-HistoMorphomics as a powerful tool for high-throughput quantification of nuclear morphology at scale and *in situ* profiling of cellular senescence, enabling high-resolution mapping of morphological and molecular features across diverse tissue types. This approach provides a foundation for mechanistic studies of senescence in aging and disease and opens new avenues for histology-based biomarker discovery.

This study demonstrates the power of high-content morphometric profiling to uncover meaningful links between cellular morphology and molecular state. A major advantage of this framework is that the biological signal can be traced to specific interpretable features that were associated with molecular or biological variables in our analyses—such as eosin texture correlation associated with SenMayo scores in endometrium, major axis length, eosin edge integrated intensity, hematoxylin radial distribution, and percent touching adjacent associated with *CCL5* expression in breast cancer stromal cells, and cell area, aspect ratio, and hematoxylin entropy associated with *CDKN1A* expression in pancreas—rather than only to abstract latent representations. Unlike patch-level representation learning approaches, sc-HistoMorphomics focuses on an explicitly defined, interpretable feature space at single-cell resolution, enabling direct biological interpretation and integration with spatially resolved molecular measurements. To more clearly position sc-HistoMorphomics within the landscape of quantitative histomorphometry and digital pathology approaches, we summarized representative prior methods according to their scale, feature representation, molecular integration, limitations relative to sc-HistoMorphomics, and intended analytical role (Supplementary Table S3).

Recent foundation models for H&E images, such as UNI^23^ and related self-supervised approaches, provide powerful patch-level representations for digital pathology tasks. However, the central contribution of sc-HistoMorphomics is different: it provides explicit, interpretable, single-cell-resolved morphology features that can be directly aligned with spatial transcriptomic measurements. We therefore view foundation-model embeddings and sc-HistoMorphomics features as complementary analytical representations, with the present study focused on the biological value of interpretable morphology alongside molecular readouts.

A key advantage of this explicit feature-based formulation is that the different morphometric categories capture complementary aspects of cellular organization. The minimal inter-category correlation among features (Figure 2) highlights the complementary nature of morphometric data across binary masks, eosin and hematoxylin staining, and neighbourhood descriptors. Such diversity allows the profiling of subtle cellular states and tissue-context effects that may be difficult to recover from more compressed or less interpretable representations. The quantitative redundancy analyses further support this interpretation. Only 2.87% of feature pairs exceeded ∣r∣>0.8, and only 1.44% exceeded ∣r∣>0.9. PCA similarly showed that 43 PCs were required to explain 90% of variance and 60 PCs were required to explain 95%, while the effective-rank estimate was 93. Together, these results indicate that the 194-feature space is not reducible to a few dominant axes, but instead captures multiple complementary morphological modes.

Because informative features often occur in partially correlated families—particularly among eosin and hematoxylin texture descriptors—feature importance is best interpreted at both the individual features and feature-family groups.

In each of our three case studies, quantitative morphology correlated with cell type and cell state, demonstrating a consistent within-sample relationship across tissues. These results also help define the discriminative resolution of H&E-derived morphology. Broad morphologically distinct cell classes were recovered more readily, whereas finer transcriptionally defined subtypes—particularly closely related stromal, lymphoid, or pancreatic subpopulations—showed more limited separability by morphology alone. Thus, sc-HistoMorphomics is best interpreted as a framework for capturing broad cell-type organization, continuous morphology-associated cell states, and morphology–molecular relationships, rather than as a standalone replacement for molecular cell typing. In the breast cancer dataset, these relationships were evident even within stromal cells, where representative morphology features distinguished *CCL5*-expressing from non-expressing stromal cells, indicating that sc-HistoMorphomics captures molecular heterogeneity within a shared cell type. Importantly, in our pancreatic example, these correlations were conserved among individuals of varying ages and disease status, suggesting that robust and interpretable imaging biomarkers that reflect complex underlying molecular profiles can be extracted from readily obtainable H&E images that are routinely collected in clinical practice.

These results show that H&E-derived morphology can reflect transcriptionally defined cell states and tissue programs, while also defining important limits of morphology-based inference. Transcriptomic measurements provide high-resolution information about molecular identity and regulatory state, whereas morphology provides an interpretable tissue-level and cell-level phenotype that can be extracted from routine histology. In this framework, spatial transcriptomics serves as a reference modality for identifying molecular states, and sc-HistoMorphomics tests whether those states are associated with reproducible and interpretable H&E-derived features. Broad morphologically distinct cell classes were recovered more readily, whereas finer transcriptionally defined subtypes showed more limited separability by morphology alone. Thus, sc-HistoMorphomics should be interpreted as a practical histology-based framework for linking tissue morphology to molecular measurements, rather than as a replacement for transcriptomic profiling or as evidence that morphology captures all molecular distinctions.

A related limitation of the endometrium Visium analysis is that morphology and gene expression are both represented at the spot level rather than at true single-cell resolution. For this reason, we summarized morphology within each spot using the mean and standard deviation of the constituent single-cell features, thereby capturing both the dominant local morphology and intra-spot heterogeneity while remaining matched to the available transcriptomic resolution. In contrast, for the Xenium datasets, where single-cell molecular data are available, we analyzed morphology directly at single-cell resolution and performed unsupervised clustering on single-cell morphology vectors. Thus, the representation used in each dataset was chosen to match the resolution of the corresponding molecular assay rather than imposed uniformly across platforms. Future work could explore richer set-based or deconvolution-style morphology representations for spot-level assays, but such approaches would go beyond the resolution directly provided by Visium.

In this study, senescence-associated cells or regions are operationally defined using transcriptomic reference labels such as the SenMayo signature and *CDKN1A*, and we then test whether they can be recognized from H&E-derived morphology. *CDKN2A* was also available in the panel, however we interpret *CDKN2A* cautiously because gene-level *CDKN2A* measurements do not necessarily distinguish the *p16*-associated signal from other products of the *CDKN2A* locus, and therefore should not be treated as an isoform-specific *p16* readout. This reflects the intended translational use case of sc-HistoMorphomics: leveraging expensive spatial transcriptomics assays to learn morphology-based signatures that can ultimately be applied to routine histology images. An important limitation is that senescence itself lacks a single universally definitive marker *in vivo*; accordingly, the morphology characterized here should be interpreted relative to the transcriptomic reference labels used in each dataset rather than as an absolute or exhaustive definition of senescence. Within that framework, our results support the feasibility of identifying transcriptomically defined senescence-associated states directly from H&E morphology.

In the endometrium Visium dataset, SenMayo-enriched spots were also enriched for stromal cells, raising the possibility that morphology–SenMayo associations could be driven by spot composition. However, the top eosin texture feature remained significantly associated with SenMayo after adjustment for stromal and epithelial enrichment, indicating that the observed signal is not solely explained by stromal identity and instead captures additional senescence-associated variation at the spot level. Thus, the endometrium result is best interpreted as morphology of senescence-enriched stromal regions, with specificity constrained by Visium’s spot-level resolution.

An additional limitation is that the observed association between eosin-derived texture features and SenMayo is correlative and does not establish causal directionality. Current understanding of cellular senescence provides several plausible mechanisms for this relationship. Senescent cells undergo broad structural, metabolic, and secretory changes, including development of the senescence-associated secretory phenotype (*SASP*), which can alter cytokine signaling, extracellular matrix composition, and local tissue organization. Senescence has also been linked to broader extracellular matrix remodeling and tissue stiffening, both of which could influence eosin staining patterns and local texture. Within this framework, eosin-associated morphology may reflect senescence-enriched states through changes in cytoplasmic composition, matrix remodeling, or tissue microenvironment. However, the current data do not distinguish whether these eosin texture changes precede, follow, or co-evolve with senescence-associated transcriptional programs. Addressing that question will require longitudinal or perturbational studies beyond the scope of the present work.^1^

Tissue sectioning is a shared limitation of both morphology-based and spatial transcriptomic analyses. Because each section captures only a two-dimensional slice through a three-dimensional tissue, nuclei and cell bodies may be partially represented or fragmented, potentially affecting apparent nuclear geometry, cytoplasmic context, neighbourhood relationships, and transcript capture. Our analyses focus primarily on relative within-sample morphology–gene expression associations, which mitigates but does not eliminate this source of variability. Future studies using serial sections, volumetric imaging, or 3D reconstruction will be important for evaluating how sectioning planes influences morphology–molecular relationships.

While our analyses primarily relied on library-size normalization, we note that normalization strategy can substantially influence gene–morphology associations. Alternative approaches such as using raw count result in quantitative differences in correlations, although the relationships between *CDKN1A* expression and morphological features remain consistent across 32 samples. This highlights both the robustness of morphology–based senescence signatures and the need for continued refinement of normalization methods for spatial transcriptomics. Because the primary analyses in this study were conducted within slides, inter-slide staining variability was not a major source of the reported morphology–gene expression associations. Within this design, the endometrium and breast cancer datasets are best interpreted as single-sample proof-of-concept applications, whereas the pancreas cohort provides the principal multi-donor validation of reproducibility in this study; future cross-slide stain normalization will be important for larger multi-cohort or multi-site applications.

Recent work has proposed more stable normalization strategies for high-plex in situ platforms such as Xenium. For example, Taylor et al.^25^ show that expression normalization using a small set of cell-type-specific, low-variation housekeeping genes may outperform global depth-based methods, particularly in settings where transcriptional output varies across cells. Although their study focuses on cancer, the concept is broadly applicable. Incorporating such approaches in future Xenium analysis may enhance comparability across samples and further strengthen morphology–gene expression relationships. These considerations underscore the value of morphological profiling as a complementary indicator of cellular states such as senescence, while acknowledging ongoing challenges in optimizing spatial transcriptomic normalization frameworks.

Segmentation quality is an important upstream source of variability for any morphology-based framework. In our comparison of StarDist and Cellpose, the most biologically informative texture-based features were highly reproducible across methods, whereas mask-derived and neighbourhood features were more sensitive to segmentation differences, particularly Zernike moment features that depend strongly on precise boundary definition. We selected StarDist as the primary segmentation method because it provides an H&E-trained model and substantially faster runtime in our workflow, while Cellpose served as a useful general-purpose comparator. We did not benchmark Mesmer directly on H&E because it is designed for multiplexed tissue imaging with separate nuclear and membrane/cytoplasmic channels. Importantly, sc-HistoMorphomics remains modular with respect to segmentation and can incorporate improved segmentation methods as they become available.

## Methods

### Experiments and data collection

The pancreas spatial transcriptomics data used in this study were generated as part of a separate study^14^. During the peer-review process, these data are available to reviewers upon request.

The endometrial datasets used in this study were generated as part of a separate study^9^. Experimental protocols and raw data are available to reviewers during the evaluation of this manuscript to ensure reproducibility.

Upon publication of the primary data study, accession numbers will be made fully public.

### Stain deconvolution

Prior to segmentation and feature extraction, H&E images were subjected to percentile-based intensity normalization to reduce the influence of extreme pixel values and staining outliers. Using the CellProfiler *split channels* module, stain separation was then performed to isolate hematoxylin- and eosin-derived intensity information before computing intensity, radial distribution, and texture features. Because the primary analyses in this study focus on within-slide morphology–gene expression relationships, inter-slide staining variability is less likely to drive the reported associations. We note, however, that additional stain normalization or batch-correction approaches could be incorporated in future cross-slide or multi-cohort analyses.

### H&E segmentation

To evaluate sensitivity to segmentation algorithm choice, we compared segmentations generated by StarDist and Cellpose on the same H&E image from the breast cancer dataset. For StarDist, we used a pretrained model designed for H&E-stained nuclei (*2D_versatile_he*)^8^, which is trained on histopathology datasets including MoNuSeg and TNBC and is better matched to the staining characteristics and nuclear morphology of histopathology images than models trained primarily on fluorescence or multiplexed imaging data. Cellpose^26^ was used as a broadly adopted general-purpose comparator, although its standard built-in models are not specifically trained for H&E histopathology images. Nuclei were matched across methods using centroid proximity with one-to-one nearest-neighbour (or mutual nearest-neighbour, depending on what you used) matching and a maximum distance threshold of 15 pixels (around 5.5 µm). For matched nuclei, corresponding morphological features were compared using Pearson and Spearman correlation, stratified by feature category (mask, eosin, hematoxylin, neighbourhood). Runtime for each segmentation method was recorded on the same image using the same computational resources. Consistent with our runtime observations, prior benchmarking on multiplexed tissue images showed that Cellpose was slower than StarDist-derived tissue models, although those comparisons were performed on TissueNet / fluorescence-like multiplexed tissue imaging rather than H&E histopathology.^27^ To approximately segment cytoplasm, we used the CellProfiler to tessellate the image plane by extending the nuclear masks either by 20 pixels or until they reached another cell’s boundary. Since there were no clear cytoplasmic boundaries in the H&E image, this segmentation was only used to calculate the number of neighbouring features.

### Nuclear morphology quantification

The 194 morphological features extracted by sc-HistoMorphomics were grouped into four major categories: mask-based shape descriptors, eosin intensity/texture features, hematoxylin intensity/texture features, and neighbourhood features. A complete list of all 194 morphological features, together with their categories, definitions, and calculation parameters, is provided in Supplementary Table S4. These features were designed to provide an interpretable, high-content representation of nuclear morphology spanning geometric shape, stain organization, and local spatial context.

#### Binary mask-based features

*Spatial Moment features*^28^: A series of weighted averages representing the shape, size, rotation and location of the object.

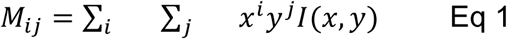

Where *i* and *j* define the order of the moment, *x* and *y* are the pixels within the image, and *I(x,y)* is the pixel intensity at *x* and *y* location where its value 1 for inside the nuclei and 0 for outside for the binary mask. Moment order matched cell profiler, *i=0,1,2* and *j=0,1,2,3*.

##### Central Moment features^28^

These features are similar to spatial moments, but they are normalized to the object’s centroid. These features are therefore not influenced by an object’s location within an image.

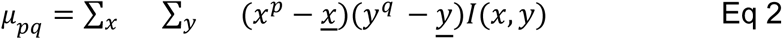

Moment order matched cell profiler, *p=0,1,2* and *q=0,1,2,3. <u>x</u>* and *y* are the center of mass of the object. The function *I(x,y)*, and *x* and *y* are the same as Spatial Moment features.

##### Normalized Moment features^28^

Similar to central moments but further normalized to be scale invariant. These moments are therefore rotation and scale invariant.

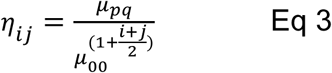

We calculate these features for *i*=0,1,2,3 and *j=0,1,2,3.* The variable *μ_pq_* is the Central Moment feature defined by eq 2.

##### Hu Moment features

Hue moments are a set of location, scale, and translation invariant features defined by Normalized Moment and central features. This means that they primarily describe the shape of the object. In this study we use *k=0,1,2,3,4,5,6*.^29^

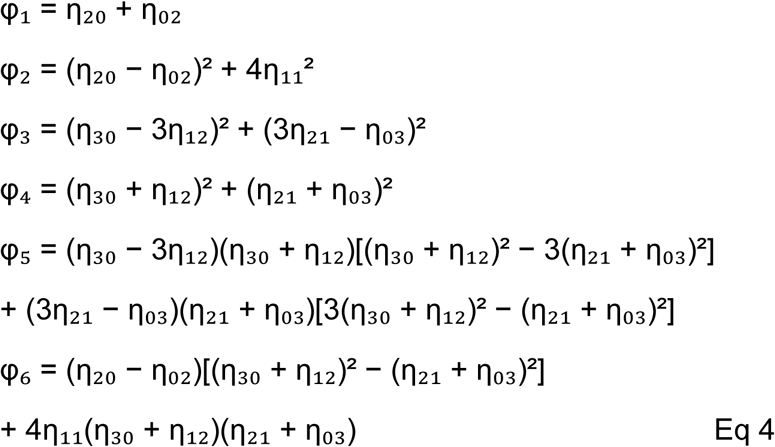

##### Inertia Tensor features^28^

These are values in the 2D inertia tensor which measure the covariance of image intensity along the image axes. These features are rotation invariant. Its components are defined as:

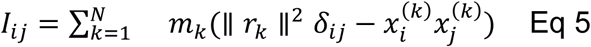

where:

- *i*, *j* ∈ {1,2,3}correspond to *x*, *y*
- *r_k_ = (x_1_^(k)^, x_2_^(k)^, x_3_^(k)^* is the position vector of mass *m_k_*,
- *δ_ij_* is the Kronecker delta.

Note that, by the definition, *I* is a symmetric tensor.

##### Inertia Tensor Eigenvalues features

These features are the relative magnitude of the eigenvalues of the tensor and represent the elongation of the object in the image^28,30^.

#### Hematoxylin- and eosin-intensity-based features

These features are divided into three categories, each capturing complementary aspects of nuclear morphology from H&E images. First, we quantified stain-intensity statistics for both eosin and hematoxylin within the nuclear mask, including measures such as mean, median, minimum, maximum, integrated intensity, and the standard deviation of pixel intensities. These features summarize global properties of staining density and reflect broad differences in chromatin condensation and cytoplasmic or extracellular matrix content. Second, we computed radial distribution features, which partition each nucleus into concentric bands and measure how stain intensity varies within each band. These features capture spatial organization of chromatin and cytoplasmic staining patterns, enabling quantification of peripheral versus central intensity enrichment. Finally, we extracted gray-level co-occurrence matrix (GLCM) features, which characterize fine-scale texture by summarizing how pixel intensities change relative to their neighbours across the nucleus. GLCM metrics such as contrast, entropy, correlation, angular second moment, and inverse difference moment quantify local intensity heterogeneity, smoothness, and structural regularity, providing sensitive measures of subtle texture patterns not captured by simpler intensity statistics. Together, these three categories form a comprehensive and interpretable set of quantitative descriptors of nuclear morphological variation^3,31,32^.

### Feature aggregation in Visium data

For Visium data, morphology was summarized at the spot level because gene expression is measured only for the spot as a whole and cannot be reliably assigned to individual cells. For each spot, nuclei whose centroids fell within the spot boundary were identified, and the mean and standard deviation of each single-cell morphology feature were computed across those nuclei, yielding 388 spot-level morphology features. The mean captures the dominant morphological state within the spot, whereas the standard deviation captures intra-spot heterogeneity among constituent nuclei. We used this representation to match the resolution of the molecular measurements and avoid introducing artificial cell-level assignments not supported by Visium.

### Tiling for parallel processing

To scale up the CellProfiler workflow to whole-slide images, we developed a pipeline for tiling large image tiles, which enabled parallel processing to speed up the shape quantification step.

To address cells that fell on tile boundaries, we gave each tile a 30-pixel buffer in all directions (substantially greater than the typical nucleus). Due to the size of the buffer, partial nuclei in one tile will fall fully within an expanded neighbouring tile. As an example, in Supplementary Figure 12 the nucleus number 3 was dropped when analyzing the red tile but was included on the blue expanded tile. Some nuclei fell into more than one tile, for example nuclei 1 and 2, which were removed based on duplicate centers of mass that were computed relative to the whole-slide image and not the tile. To avoid pulling the whole image into the memory and parallelization, we used the dask arrays (which allow large images to be broken into manageable chunks and processed in parallel without loading the entire slide into memory) to chunk the whole-slide image and use the map_overlap function to expand the tiles and apply the CellProfiler pipeline to each chunk. We later adjusted the cell center of mass information in the feature data.

### Feature redundancy and dimensionality analysis

Feature redundancy and dimensionality were evaluated using pairwise Pearson correlations, effective-rank analysis, and principal component analysis. Pairwise Pearson correlations were calculated among z-scored morphology features. To quantify redundancy, the upper triangle of the pairwise correlation matrix was extracted, excluding the diagonal, and the fraction of feature pairs exceeding high absolute-correlation thresholds, including ∣r∣>0.8|r| and ∣r∣>0.9, was calculated. Pairwise correlations were also summarized by feature-category pair.

PCA was performed on the z-scored morphology feature matrix. The proportion of variance explained by each principal component was calculated from the PCA eigenvalue spectrum, and cumulative variance was used to determine the number of components required to explain 90% and 95% of the total variance.

To quantify the intrinsic dimensionality of the feature space, we computed the effective rank of each tissue-specific data matrix^10^. Before analysis, each feature was standardized to zero mean and unit variance. For a given matrix *X*, we calculated its singular value decomposition *X* = *U* ∑ *V^T^* and obtained the non-negative singular values {*σ*}. We then converted these values into a normalized distribution 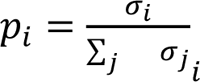.

The effective rank was defined as the exponential of the spectral entropy,

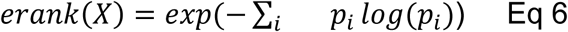

which yields a continuous estimate of the number of dominant orthogonal directions contributing to the variance structure of the data. This measure ranges from 1 (variance concentrated in a single direction) to 194, the number of features. We report the effective rank separately for each tissue to quantify how uniformly variance is distributed across singular directions and to assess redundancy relative to the full feature space.

### Calculating Enrichment score

To calculate the enrichment score we used the Giotto package in R. This package takes marker genes of interest as input and calculates the enrichment score for that gene set in each spot. As senescence marker gene sets, we used the SenMayo marker gene signature introduced in Saul et al^33^ and listed in Table S1. As cell type markers, we used gene sets reported in Tan et al.^34^ for the different cell types in the endometrium data set. These gene sets are also listed in Table S2.

### Random forest prediction of SenMayo score

To build the model to predict SenMayo enrichment score from morphology features in the endometrium data, we used a random forest regression model from scikit-learn package in python. We split the spots into training (75%) and testing (25%) data sets. The maximum tree depth used was 20 and the number of features used to consider for the best split was 2.

### Within-stromal analysis of *CCL5*-associated morphology in breast cancer

For the breast cancer Xenium dataset, we evaluated *CCL5*–morphology associations within stromal cells only to control for broad cell-type differences. Stromal cells were divided into *CCL5*-expressing and non-expressing groups based on whether normalized *CCL5* expression exceeded zero. Representative morphology features were compared between the two groups using two-sided Wilcoxon rank-sum tests. In addition, continuous associations between morphology features and *CCL5* expression within stromal cells were quantified using Spearman correlation.

Unsupervised clustering of single-cell morphology features

For the Xenium breast cancer and pancreas datasets, we first z-scored single-cell morphology feature vectors and then performed unsupervised clustering using k-means, with k = 3 for the breast cancer dataset and k = 4 for the pancreas dataset. These values were chosen to capture the dominant broad cell types annotated in each tissue. Resulting morphology-based clusters were compared to post hoc with expression-derived cell-type annotations.

## Code availability

The sc-HistoMorphomics pipeline is publicly available at the <u>Sc-HistoMorphomics</u> <u>GitHub repository</u>. The repository includes documentation, example workflows for nuclear segmentation and morphological feature quantification using the breast cancer dataset, and scripts required to reproduce the main figures presented in this study.

## Data availability

The breast cancer dataset analyzed in this study is publicly available from its original source through the 10x Genomics Xenium Human Breast Dataset.

## Contributions

E.A., B.S.W., S.L., and J.M.M. conceived the study. E.A., Y.Z., S.D., D.B., B.S.W., and J.M.M. performed data curation. E.A., B.S.W., and J.M.M. performed data analysis. E.A. performed data visualization. S.D., D.B., D.U., W.F.F., J.H.C., E.C., B.S.W., N.R., P.R., R.K., and J.M.M. provided resources. B.S.W., N.R., P.R., R.K., S.L., and J.M.M. supervised the study. N.R., P.R., J.H.C., E.C., B.S.W., S.L., and J.M.M. acquired funding. E.A., B.S.W., and J.M.M. wrote the original draft of the manuscript. All authors reviewed and edited the manuscript.

## Acknowledgements

This work is supported by the NIH Common Fund, through the Office of Strategic Coordination/Office of the NIH Director under awards U54AG079753 (JAX-Sen) and U54AG075941 (KAPP-Sen) as part of the SenNet Consortium.

