## Supplementary Figures for "Sc-HistoMorphomics: A General Framework for High-throughput, Multimodal Profiling of Nuclear Morphology in Tissues"

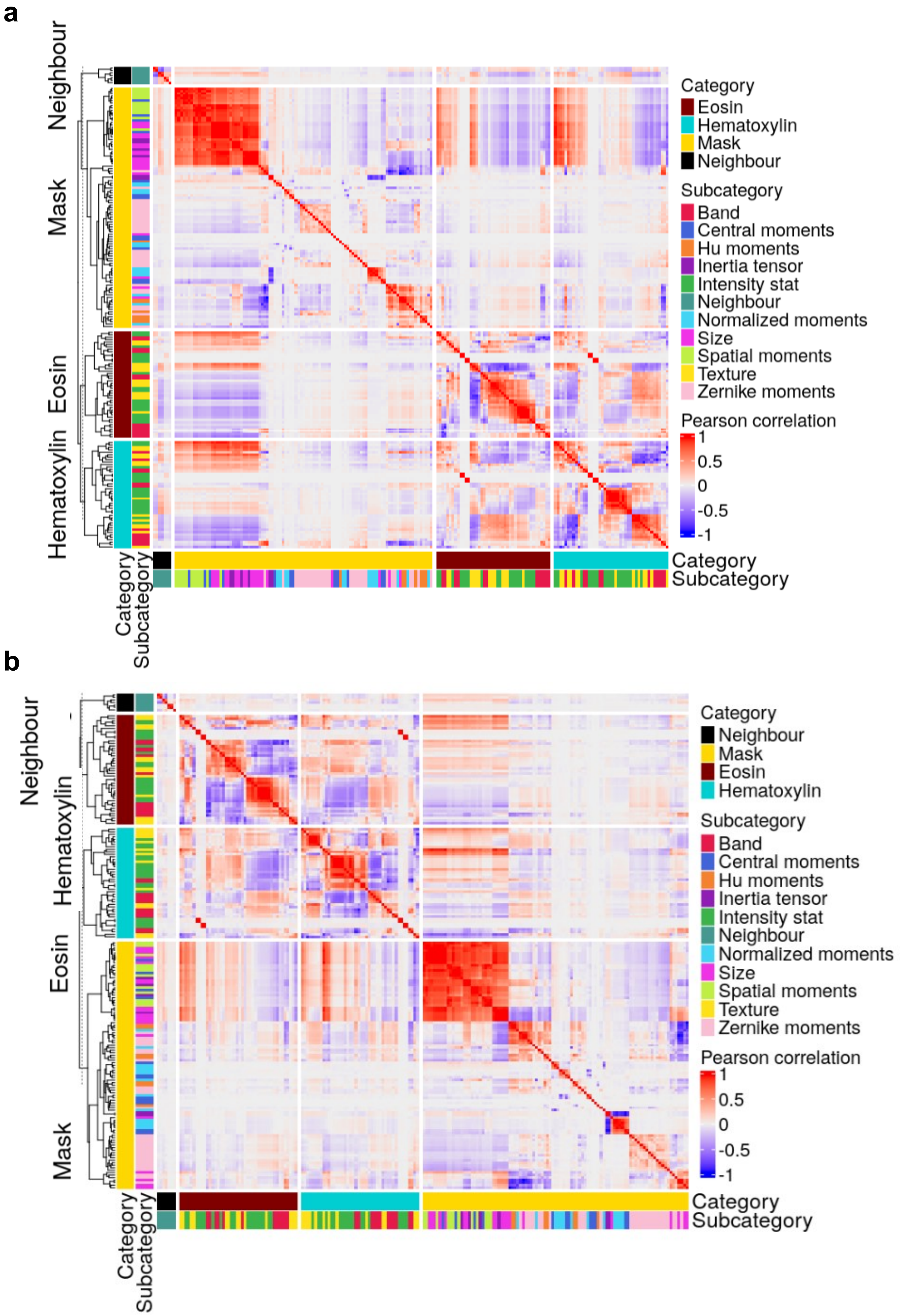


**Supplementary Figure S1. The four major morphological categories are largely independent for Breast cancer and pancreas tissue.** Feature correlations in a) breast cancer, and b) pancreas tissue is very similar. Both have many redundant features.

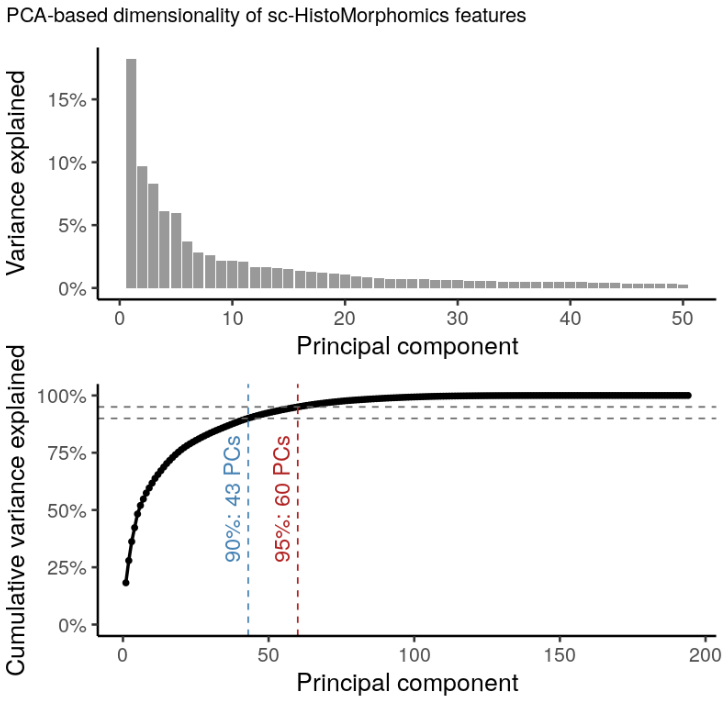


**Supplementary Figure S2. PCA-based dimensionality of sc-HistoMorphomics features.** Scree plot and cumulative variance curve from PCA of the z-scored morphology feature matrix. Variance was distributed across many components, with 43 PCs required to explain 90% of variance and 60 PCs required to explain 95%, supporting a high-dimensional, partially non-redundant feature space.

a

b

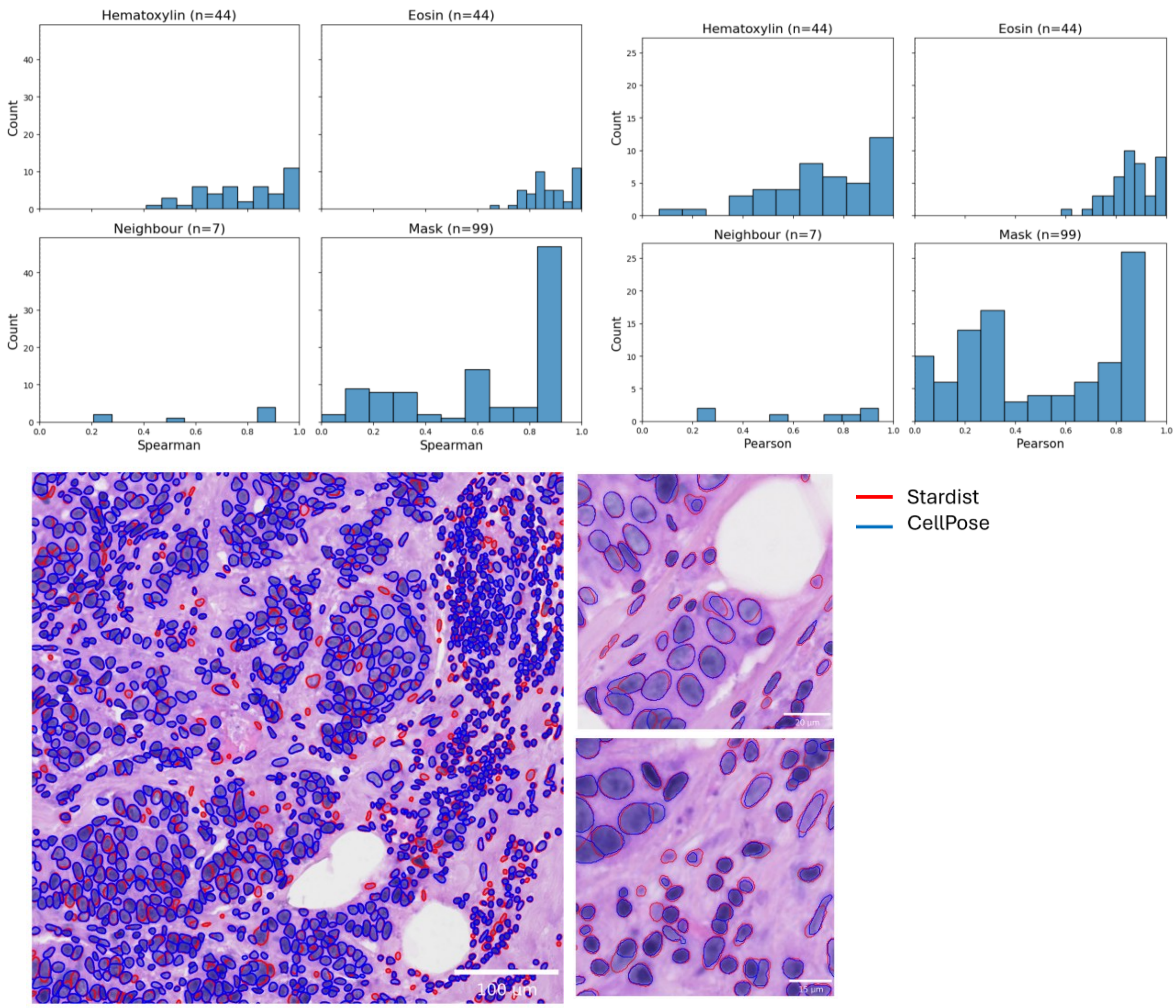


**Supplementary Figure S3. Segmentation-method robustness of sc-HistoMorphomics features.** a) Feature-wise agreement between morphology measurements extracted from nuclei segmented using two independent algorithms, StarDist and Cellpose, in the same breast cancer H&E image. Nuclei were matched across segmentation methods, and corresponding feature values were compared using Spearman correlation coefficients and Pearson correlation coefficients, shown separately by feature category. Hematoxylin and eosin texture features each included 44 features and showed the strongest cross-method agreement, with most correlations concentrated at high values. Mask-derived features included 99 features and showed a broader distribution of correlations, consistent with the greater sensitivity of boundary-dependent shape descriptors to small differences in nuclear segmentation. Neighborhood features included 7 features and showed the greatest variability, likely reflecting their dependence on local spatial relationships and the smaller number of features in this category. Overall, texture-based features were highly reproducible across segmentation methods, whereas mask-derived and neighborhood features were more sensitive to segmentation differences. b) Representative nuclear segmentation results obtained with StarDist. Segmentation boundaries (red) are overlaid on the original image to demonstrate segmentation accuracy across nuclei with varying size, density, and morphology. Cellpose segmentations (blue) are shown for comparison.

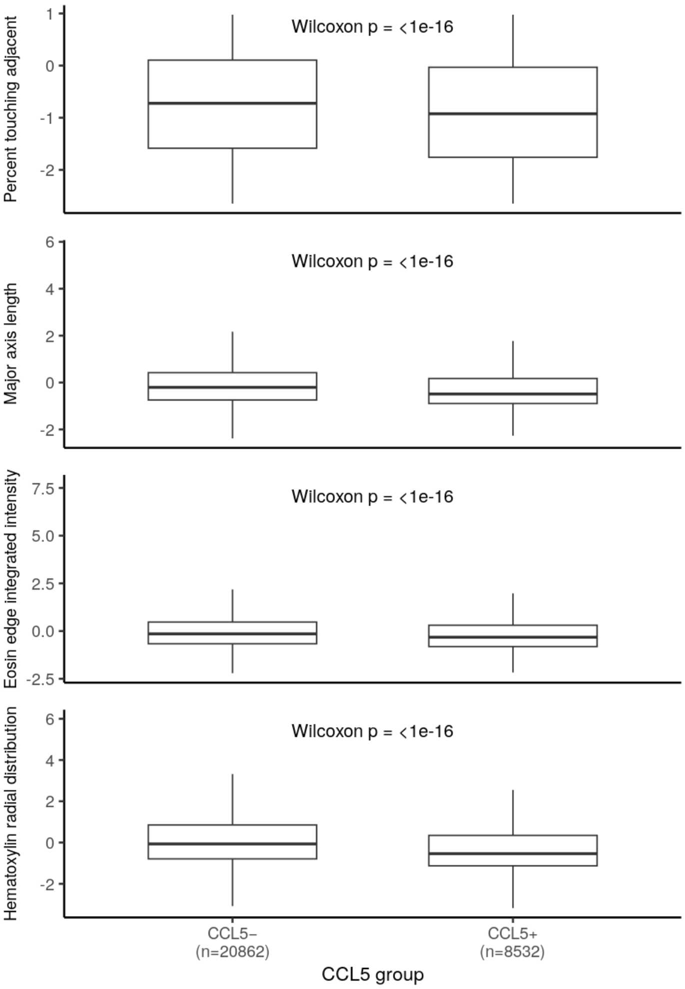


**Supplementary Figure S4.** Morphology distinguishes CCL5-expressing and non-expressing stromal cells in breast cancer tissue. Boxplots compare representative morphology features between CCL5-negative and CCL5-positive stromal cells, with sample sizes and Wilcoxon rank-sum p-values shown. Binary expressor/non-expressor comparisons and continuous expression correlations capture related but distinct aspects of molecular heterogeneity and are therefore not necessarily expected to yield identical directionality for every feature.

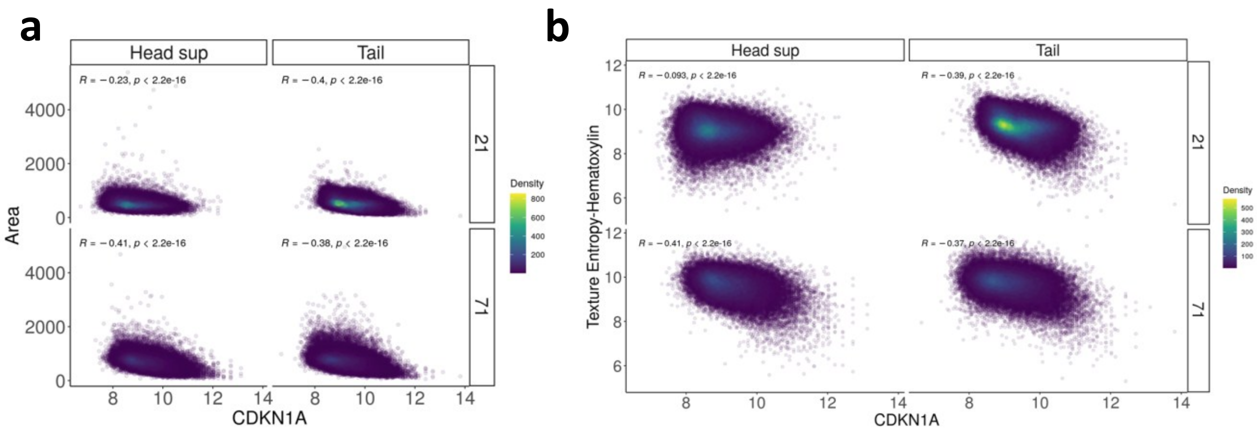


**Supplementary Figure S5. The correlation between morphological features and *CDKN1A* is consistently observed across samples.** *CDKN1A* expression is negatively correlated with a) area of nuclei and b) texture entropy feature of hematoxylin stain in the “head” and “tail” regions of two patients.

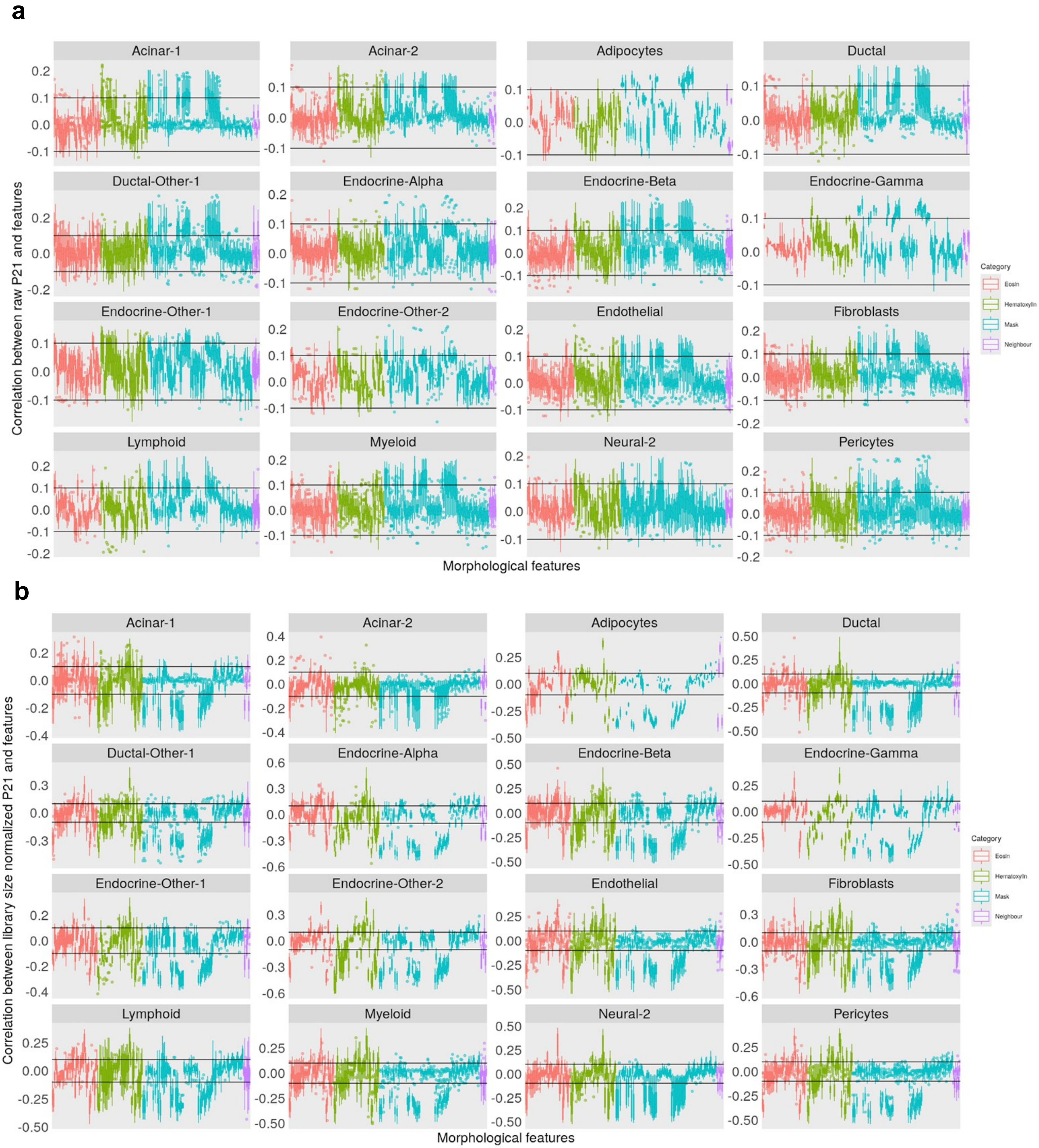


**Supplementary Figure S6. There is repeatable positive correlation between morphological features and *CDKN1A* expression regardless of normalization method.** a) Correlation between features and raw *CDKN1A* expression for different cell types, in 32 different pancreas samples. b) Correlation between features and library size normalized *CDKN1A* expression for different cell types, in 32 different pancreas samples. Each datapoint for each boxplot is the correlation between a morphological feature and P21 expression per cell type per sample.

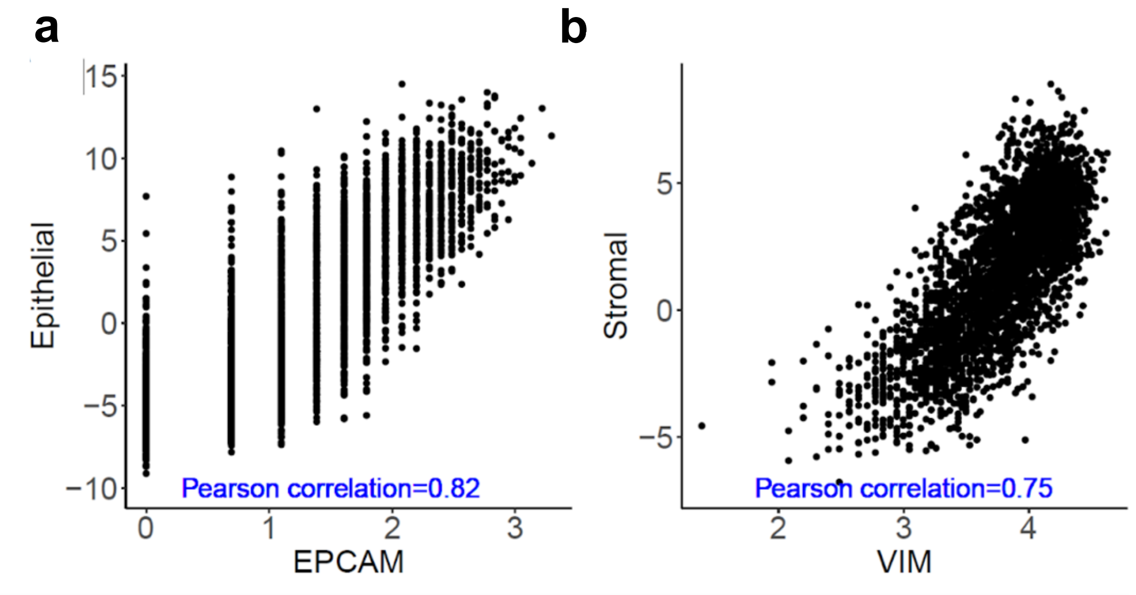


**Supplementary Figure S7. Cell type enrichment scores are significantly correlated with the well-known markers for each cell type.** a) Epithelial enrichment score is correlated with *EPCAM*. b) Stromal enrichment score is correlated with *VIM*.

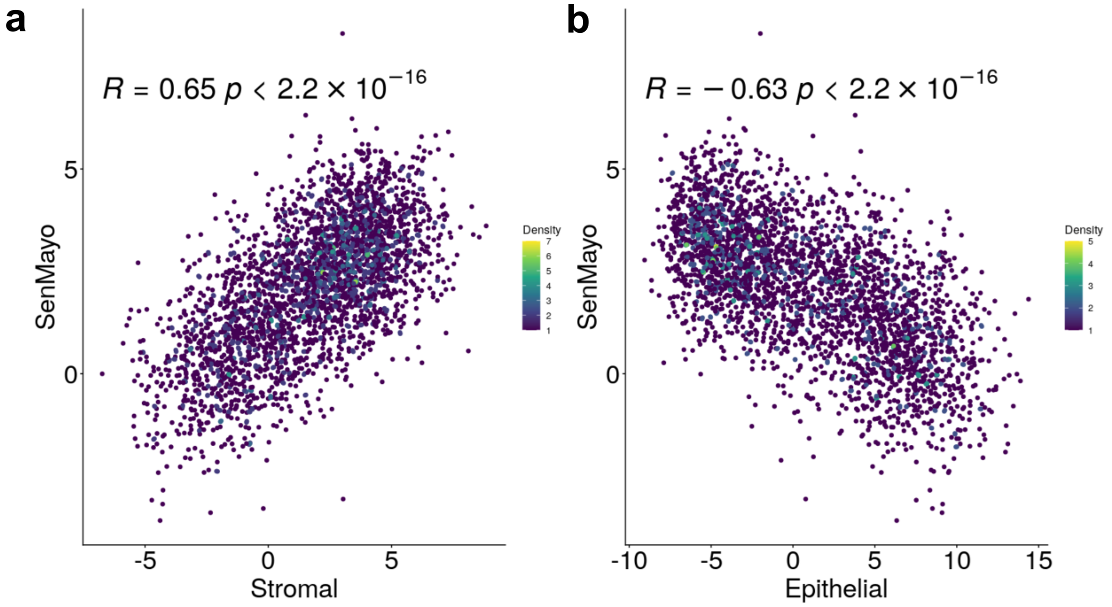


**Supplementary Figure S8. SenMayo score, epithelial score and stroma score are correlated.** Spots with a high SenMayo score were enriched for stromal cells and depleted for epithelial cells in the endometrium. This shows that stromal cells are enriched with senescence cells.

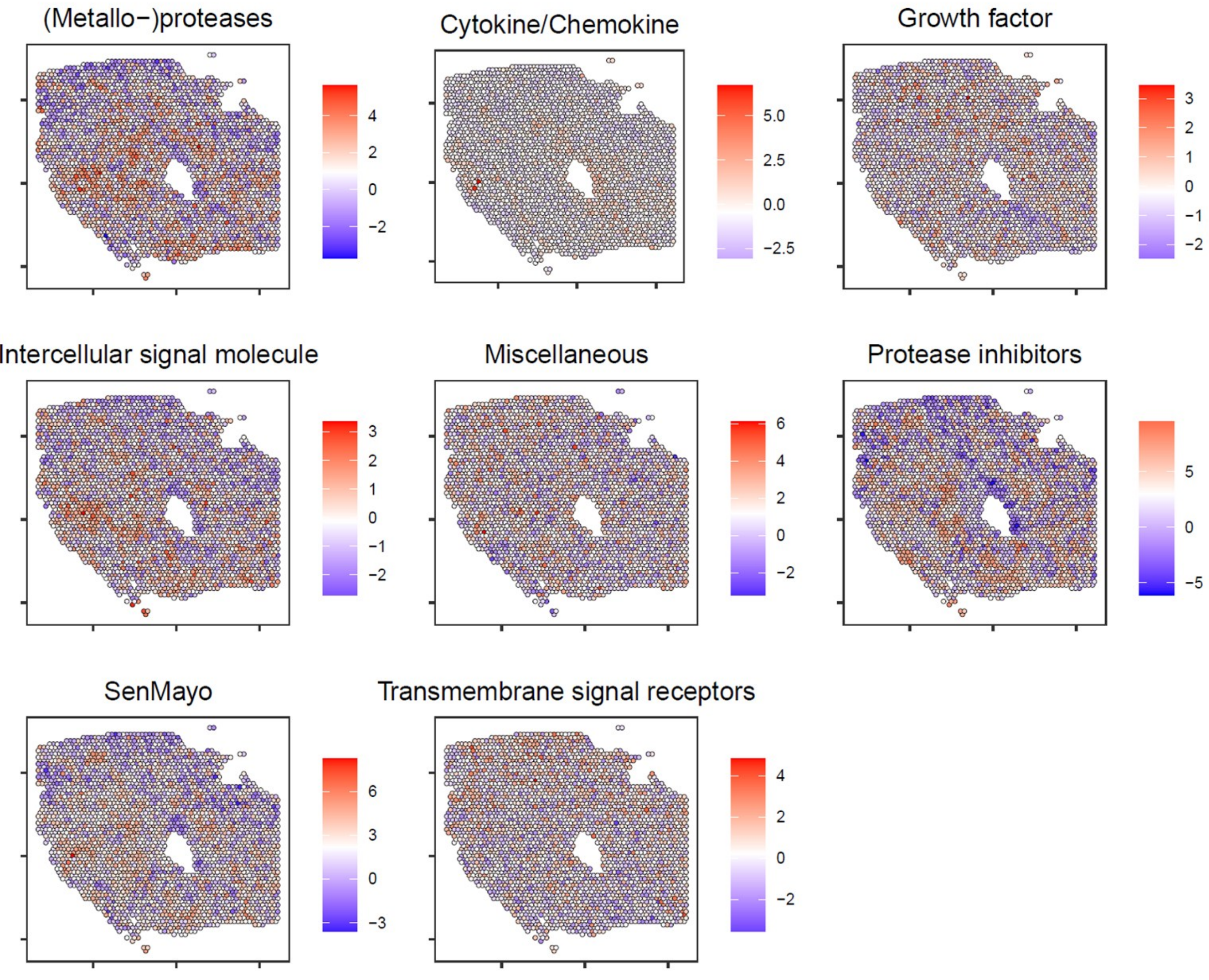


**Supplementary Figure S9. Senescent cells are dispersed across the tissue.** The enrichment score for different senescence pathways is very heterogeneous across the tissue, especially for protease inhibitors, metallo-proteases, and SenMayo pathways.

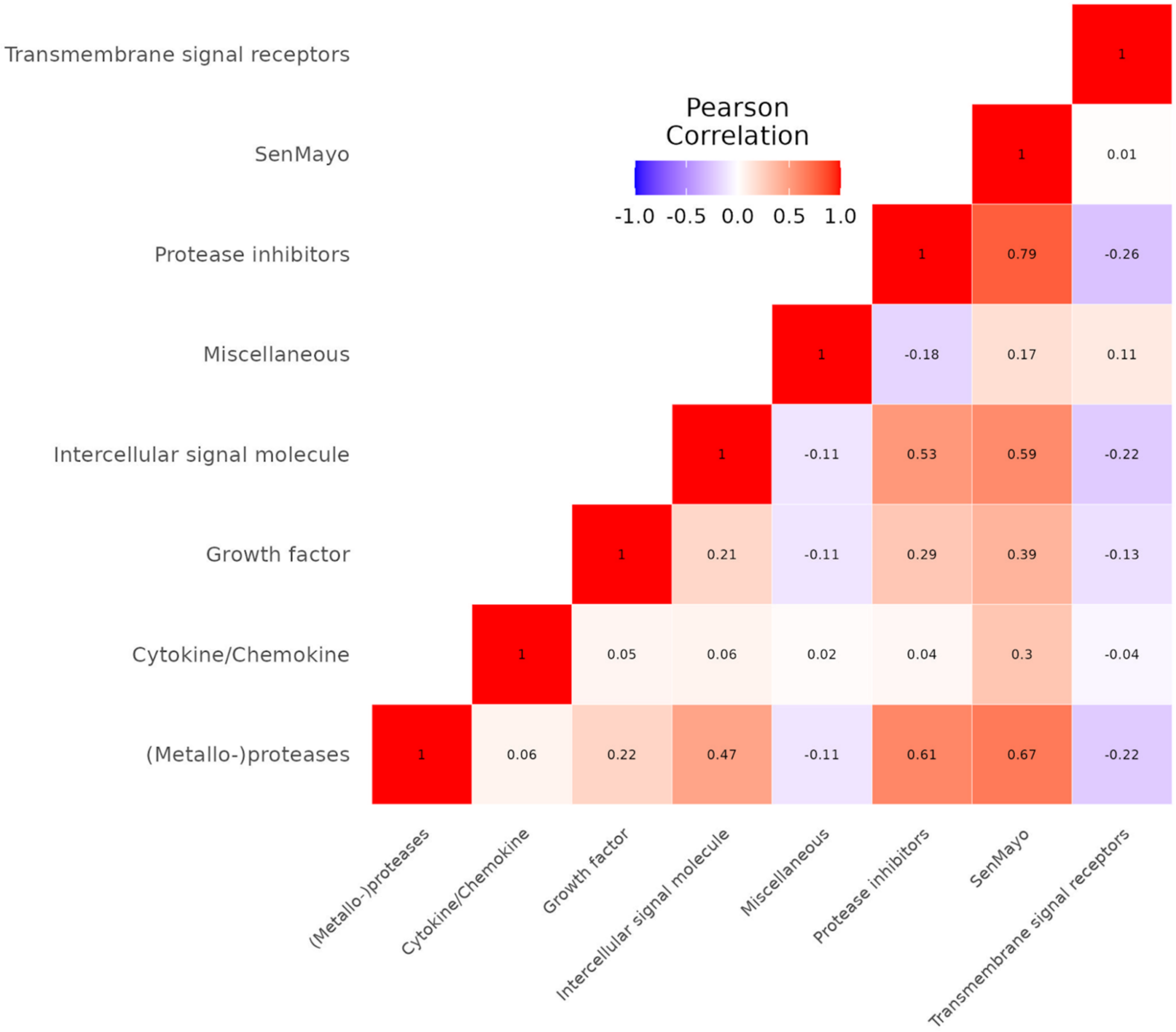


**Supplementary Figure S10. The Pearson correlation between different SenMayo subcategories show moderate positive correlations.** This shows that these subcategories capture related but biologically distinct senescence-associated processes since they contain non-overlapping genes.

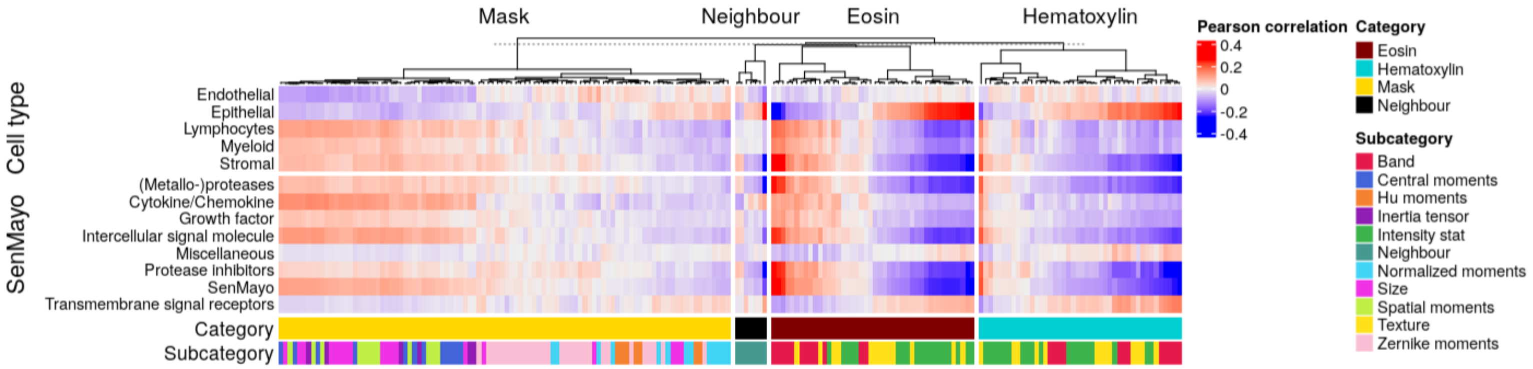


**Supplementary Figure S11. Standard deviation of eosin features within each spot are morphological markers of senescence in the endometrium.** Pearson correlation between standard deviation of morphological features within a spot and either spot-level SenMayo scores or cell type enrichments. Eosin markers show high correlation with SenMayo score.

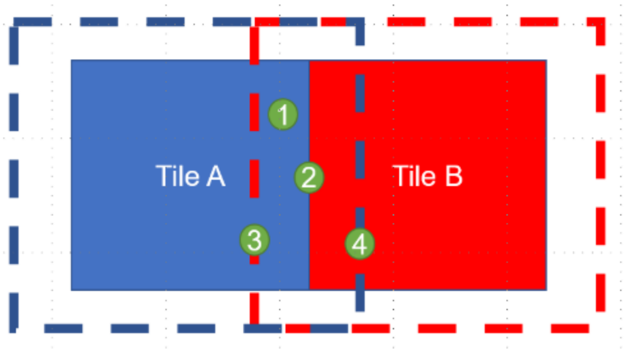


**Supplementary Figure S12. Workflow runs the tissue in tiles.** Each tile is expanded to include 30-pixels from each neighbouring tile. Nuclei that fall on the boundary of the tile are dropped since their morphological features will not be accurate. Nucleus number 3 is dropped when analyzing the red tile but is included on the blue expanded tile. When nuclei fall into more than one tile (nuclei 1 and 2), only one of them will be used in downstream analysis.
